# A genome-wide CRISPR screen identifies interactors of the autophagy pathway as conserved coronavirus targets

**DOI:** 10.1101/2021.02.24.432634

**Authors:** Annika Kratzel, Jenna N. Kelly, Yannick Brueggemann, Jasmine Portmann, Philip V’kovski, Daniel Todt, Nadine Ebert, Eike Steinmann, Ronald Dijkman, Gert Zimmer, Stephanie Pfaender, Volker Thiel

## Abstract

Over the past 20 years, the emergence of three highly pathogenic coronaviruses (CoV) – SARS-CoV, MERS-CoV, and most recently SARS-CoV-2 – has shown that CoVs pose a serious risk to human health and highlighted the importance of developing effective therapies against them. Similar to other viruses, CoVs are dependent on host factors for their survival and replication. We hypothesized that evolutionarily distinct CoVs may exploit similar host factors and pathways to support their replication cycle. Here, we conducted two independent genome-wide CRISPR/Cas9 knockout screens to identify pan-CoV host factors required for the replication of both endemic and emerging CoVs, including the novel CoV SARS-CoV-2. Strikingly, we found that several autophagy-related genes, including the immunophilin FKBP8, TMEM41B, and MINAR1, were common host factors required for CoV replication. Importantly, inhibition of the immunophilin family with the compounds Tacrolimus, Cyclosporin A, and the non-immunosuppressive derivative Alisporivir, resulted in dose-dependent inhibition of CoV replication in primary human nasal epithelial cell cultures that resemble the natural site of virus replication. Overall, we identified host factors that are crucial for CoV replication and demonstrate that these factors constitute potential targets for therapeutic intervention by clinically approved drugs.

## Introduction

Coronaviruses (CoVs) are positive-sense single-stranded enveloped RNA viruses with a broad host tropism and in case of the three highly pathogenic zoonotic CoVs the ability to cross species barriers and infect humans. Since 1960, seven human CoVs (HCoVs) with a suspected zoonotic origin in bats, mice, or domestic animals have been identified, including four seasonally circulating well-established human pathogens (HCoV-229E, HCoV-OC43, HCoV-NL63, and HCoV-HKU1) that usually cause mild symptoms like the common cold and/or diarrhea in immunocompetent patients^1,2,3,4^. HCoV infections have therefore generally been considered harmless; however, the relatively recent emergence of three highly pathogenic HCoVs, which infect the upper and also lower respiratory tract and cause severe disease in humans, has demonstrated that HCoVs can impact human health. Between 2002 and 2003 the highly pathogenic Severe Acute Respiratory Syndrome (SARS) coronavirus was responsible for an outbreak of severe viral pneumonia causing disease in over 8,000 patients^5^. Moreover, the emergence of Middle East Respiratory Syndrome (MERS) in 2012 marked the second occurrence of a highly pathogenic CoV in humans and has persistently caused endemics in the Middle East via zoonotic transmissions from dromedary camels and nosocomial outbreaks^6,7,8^. Recently, the newly emerged SARS-CoV-2, the causative agent of coronavirus disease 2019 (COVID-19), continues to create an imminent threat to global health, with more than 100 Mio individuals currently infected in > 200 countries and more than 2 Mio fatalities (February 8^th^ 2021) (Johns Hopkins Coronavirus Resource Center).

The lack of specific pharmaceutical intervention options and/or prevention measures against HCoVs, as well as the ongoing difficulties containing the rapid global spread of the SARS-CoV-2, have intensified in the current pandemic and new therapies are urgently needed. CoVs are obligate intracellular pathogens and thus rely on selected host proteins, termed host dependency factors (HDFs), to achieve virus entry, replication, and release. The identification of HDFs is therefore of importance for the understanding of essential host-virus interactions required for successful viral replication and provide a framework to guide development of new pharmacological strategies for the treatment of CoV infections, including the disease COVID-19 and future emerging CoVs. Coronaviruses encode a spike surface glycoprotein, which enables specific binding to a host-cell receptor to mediate viral entry. Known host receptors include dipeptidyl peptidase 4 (DPP4) for MERS-CoV, human aminopeptidase N (ANPEP) for HCoV-229E, and angiotensin-converting enzyme 2 (ACE2) for SARS-CoV and SARS-CoV2^9,10,11,12^. Cleavage of the spike protein by cellular proteases, such as TMPRSS2, cathepsin L, and/or furin facilitates membrane fusion followed by release of the viral genome into the cellular cytoplasm for replication^13^. One hallmark that occurs in host cells during replication of positive-stranded RNA viruses is the extensive remodeling of host endomembranes that results in coronavirus infection in the formation of both double-membrane vesicles (DMVs) and convoluted membranes (CM) to which the viral replication and transcription complexes are targeted^14,15,16^. However, the host factors that are required for the formation of these structures remain elusive. Newly synthesized viral RNA is assembled to viral particles at the ER-Golgi intermediate compartment (ERGIC) and trafficked to the Golgi for post-translational modifications^17^. While only little is known on how HCoVs exit from infected cells, recent work found that the β-CoVs MHV and SARS-CoV egress from cells via a lysosome-based pathway^18^.

To identify HDFs essential for CoV infection, we performed two independent genome-wide loss-of-function CRISPR screens with MERS-CoV, a highly pathogenic CoV, and HCoV-229E, an endemic CoV that causes mild respiratory symptoms in humans. We sought to uncover HDFs required for infection by a wide range of CoVs, including highly pathogenic CoVs with pandemic potential. Our results revealed that a number of autophagy-related genes, including FK506 binding protein 8 (FKBP8), transmembrane protein 41B (TMEM41B), vacuole membrane protein 1 (VMP1), and Membrane Integral NOTCH2 Associated Receptor 1 (MINAR1), were among the top hits for both CoV screens, suggesting that host factors involved in autophagy may also be required for CoV replication. Importantly, we found that perturbation of FKBP8 and other members of the immunophilin family by clinically approved and well-tolerated drugs, but not inhibition of late cellular autophagy, inhibited CoV infection in a dose-dependent manner. Overall, the genes and pathways identified in our CoV screens expand the current repertoire of essential HDFs required for CoV replication that can be exploited to identify novel therapeutic targets for host-directed therapies against both existing and future emerging CoVs.

## Results

### Two independent genome-wide CRISPR/Cas9 knockout screens reveal CoV host dependency factors

We performed two independent genome-wide loss-of-function CRISPR screens with MERS-CoV and HCoV-229E to uncover unknown HDFs required for CoV replication. To conduct these CRISPR screens, we employed the well-established Human GeCKOv2 genome-wide library, which includes 65,386 unique single guide RNAs (sgRNAs) targeting 19,052 protein-coding genes^19^. As a screening platform, we selected human hepatoma Huh7 cells for several reasons. First, Huh7 cells endogenously express DPP4 and ANPEP/CD13, the host cell receptors for MERS-CoV and HCoV-229E, respectively^9,10^. Thus, Huh7 cells are susceptible to infection with both viruses. Second, both MERS-CoV and HCoV-229E induce cytopathic effects in Huh7 cells following infection, which allows for rapid selection of CRISPR knockout-mediated non-susceptible cells. Finally, several recent studies have also selected Huh7 cells for the CRISPR-based screening of other CoVs, including the novel, highly pathogenic SARS-CoV-2 virus^20–22^.

Genome-wide CRISPR/Cas9 knockout screens were performed by transducing Huh7 cells with the Human GeCKOv2 library, selecting for library-transduced cells with puromycin, followed by infection with either MERS-CoV (37°C, MOI 0.05) or HCoV-229E (33°C, MOI 0.1). Surviving cells were harvested 14 days post infection, genomic DNA was extracted, and sgRNA abundance was quantified using amplicon-based Illumina next-generation sequencing (NGS) (Figure 1A). Technical performance was evaluated using a number of quality control metrics, including an area under the curve (AUC) analysis of all sgRNAs found in samples from each screen. AUC analysis confirmed that library representation was diverse and properly maintained in uninfected samples from both screens. As expected, AUC analysis also revealed a much greater level of sgRNA guide dropout following infection with either MERS-CoV or HCoV-229E (Figure S1A). Pairwise correlation analysis showed that biological replicates from each screen clustered together and shared a high correlation coefficient (Figure S1B).

**Figure 1:**
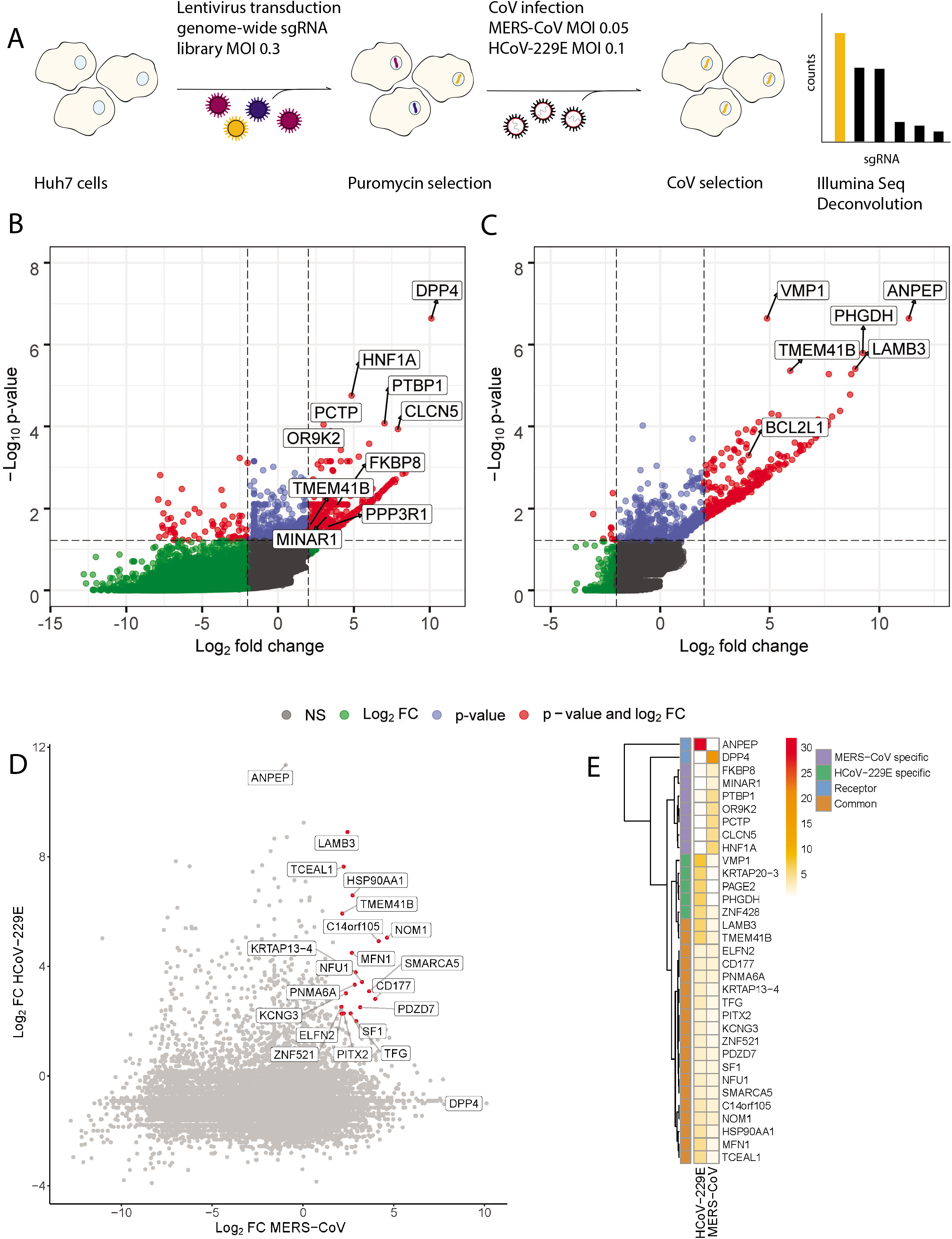
MERS-CoV and HCoV-229E genome-wide CRISPR/Cas9-mediated knockout screens. (A) Native Huh7 cells were transduced with the GeCKOv2 lentiviral genome-wide CRISPR library, ensuring a coverage of ~500 cells per sgRNA. Transduced cells were selected and then infected with either MERS-CoV or HCoV-229E at indicated MOIs and temperatures. Surviving cells were harvested and prepared for deep sequencing. Deconvolution identified both virus-specific and pan-coronavirus host dependency factors (HDFs). (B) Volcano plot showing the log fold change (Log2 FC) and log_10_ p-value for each gene in the MERS-CoV CRISPR screen. Genes with a FC ≥ 2 and p_-_value < 0.05 are highlighted in red. Selected top genes are annotated in the plot, including the MERS-CoV receptor (DPP4) and the 5 most highly ranked genes in the MERS-CoV screen. (C) Volcano plot showing the Log2 FC and log_10_ p-value for each gene in the HCoV-229E CRISPR screen. Genes with a FC ≥ 2 and p-value < 0.05 are highlighted in red. Selected top genes are annotated, including the HCoV-229E receptor (ANPEP) and the 5 most highly ranked genes in the HCoV-229E screen. (D) Pairwise comparison of enriched genes in the HCoV-229E and MERS-CoV CRISPR screens. Dotted lines indicate a Log2 FC ≥ 2. Genes with a Log2 FC ≥ 2 and p-value < 0.05 in both screens are highlighted in red and annotated. (E) Heatmap comparing the log RRA p-values for selected top virus-specific and common hits in both CoV screens. CoV receptors (DPP4 and ANPEP) are demarcated by the blue boxes, MERS-CoV specific genes by the purple boxes, and HCoV-229E specific genes by the green boxes. Common significantly enriched genes, which are also annotated in Figure 2D, are demarcated by the orange boxes. Heatmap clustering was performed using the complete linkage method and Euclidean distance.

Using the MAGeCK pipeline^23^, we performed paired analyses on uninfected and infected samples from each screen and computed gene-level scores to identify genes that were significantly enriched in our MERS-CoV and HCoV-229E infected samples. Overall, we identified 1149 genes in the MERS-CoV screen and 517 genes in the HCoV-229E screen that had significant robust rank aggregation (RRA) enrichment (p < 0.05) using the gene log fold change (LFC) alpha median method. RRA analysis using the second-best LFC method identified 989 significantly enriched genes in the MERS-CoV screen and 332 significantly genes in the HCoV-229E screen. To prioritize genes and generate a more robust dataset, we focused on genes identified as significantly enriched using both methods (RRA p-value < 0.05) with a LFC of ≥ 2 (Figures S1C and S1D). In total, 944 genes from the MERS-CoV screen and 332 genes from the HCoV-229E screen met these criteria, including 19 genes that were identified by both methods in both screens (Figures 1B, 1D, and S1D). Top scoring genes from both screens are shown in Figure 1E, including several virus-specific genes as well as the 19 aforementioned common genes. Coronaviruses encode a spike surface glycoprotein, which enables specific binding to a host-cell receptor to mediate viral entry. Known host receptors include dipeptidyl peptidase 4 (DPP4) for MERS-CoV, human aminopeptidase N (ANPEP) for HCoV-229E, and angiotensin-converting enzyme 2 (ACE2) for SARS-CoV and SARS-CoV2 ^24,10,11,12^, Subsequent cleavage of the spike protein by cellular proteases, such as TMPRSS2, cathepsin L, and/or furin enables membrane fusion followed by release of the viral genome into the cellular cytoplasm for replication^13^. Importantly, in the MERS-CoV screen, the DPP4/CD26 host cell receptor was identified as the top scoring gene, whereas in the HCoV-229E screen, the top scoring gene was ANPEP/CD13. Moreover, the known DPP4 transcription factor HNFA1 was ranked second in the MERS-CoV screen, demonstrating the robustness of the screen.

To identify and compare host cell biological processes that may be required for CoV replication, we next performed Gene Ontology (GO) enrichment analysis on each screen using the enriched genes identified above. This analysis uncovered multiple biological processes (BP) that were significantly enriched in both CoV screens, many of which clustered together into 7 overarching biological themes (Figure 2A). Next, we calculated the semantic similarity among the 636 unique GO terms (BP) that were identified as significantly enriched in one or both screens (p-value < 0.05; Table 2). Hierarchical clustering was then used to group similar GO terms together and a representative term for each group was selected based on scores assigned to each term. The latter analysis led to the identification of 44 conserved representative GO terms and 51 virus-specific representative GO terms (Figure S2A). Representative GO terms found in both MERS-CoV and HCoV-229E screens included a number of immune-related terms as well as terms related to the regulation of phosphorylation, kinase activity, autophagy, and lipid transport. Several specific GO terms were also significantly enriched in both screens, including neutrophil-mediated immunity, regulation of protein dephosphorylation, and regulation of the c-Jun N-terminal kinase (JNK) cascade (Figure S2B). GO terms specific to our MERS-CoV screen included regulation of exit from mitosis, protein glycosylation, and syncytium formation via plasma membrane fusion. In contrast, GO terms specific to HCoV-229E included regulation of coagulation and nitric oxide biosynthesis (Figure S2A).

**Figure 2:**
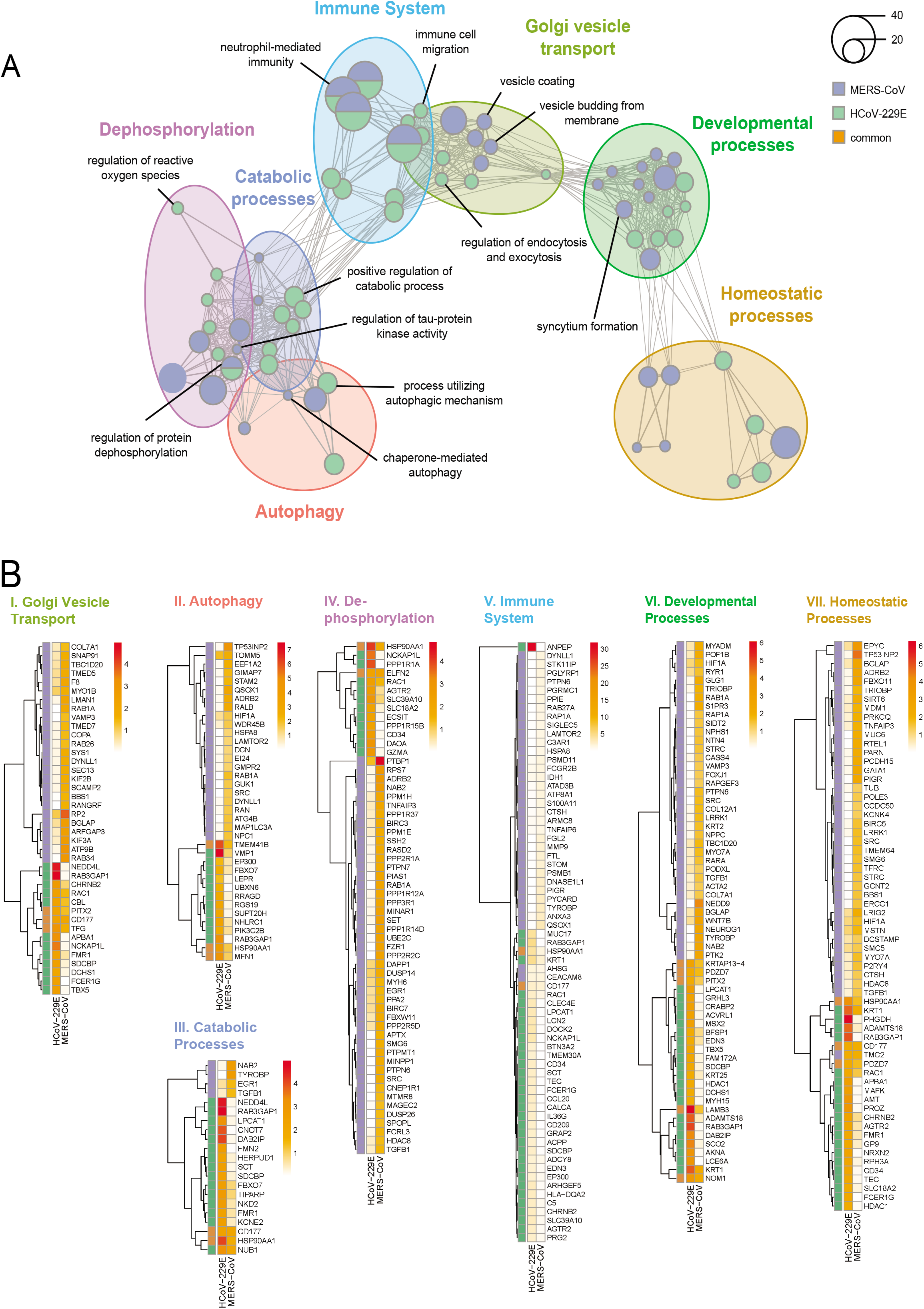
Enrichment analysis uncovers host biological networks crucial for CoV replication. (A) Enrichment map summarizing major host biological networks co-opted by CoVs during infection. Gene Ontology (GO) enrichment analysis was performed using hits from both MERS-CoV and HCoV-229E CRISPR screens and filtered to contain conserved representative GO terms and genes. Each node represents an individual GO term and nodes that are functionally related cluster together into a larger network. Node size reflects number of significantly enriched genes in the node and color indicates the CoV screen for which the node was significant. A complete list of significant GO terms can be found in Table 2. **(B)** Heatmaps of individual biological clusters displayed in **(A)**. Heatmaps contain significantly enriched genes from both CoV screens that were associated with significantly enriched GO terms found within the individual biological clusters in (A). Colored panels on the left-hand side of heatmaps show which CoV screen contained specific enriched genes (purple: MERS-CoV, green: HCoV-229E, and orange: enriched in both CoV screens). Colors in each legend represent the log RRA p-values for each gene in each CoV screen. Heatmap clustering was performed using the complete linkage method and Euclidean distance.

To establish which pathways and/or processes may be particularly important for CoV replication, we next focused on conserved representative GO terms that included one or more of the 19 genes that were significantly enriched in both of our CoV screens (Figures 1D and 1E). The resulting 70 unique GO terms and their relationships to each other are the terms illustrated in Figure 2A. The 7 prominent biological themes these 70 terms clustered into are also shown and include autophagy, immunity, dephosphorylation, Golgi vesicle transport, catabolic processes, homeostatic processes, and developmental processes. To examine each biological cluster in more detail, we constructed cluster-specific heatmaps showing all enriched genes from both CoV screens associated with that cluster (Figure 2B). Furthermore, for each cluster we inspected the network of functionally related GO terms that comprise the cluster (Figure S3A-G). Overall, our results indicate the involvement of diverse biological processes in both, MERS-CoV and HCoV-229E replication cycle.

### Regulators of the autophagy pathway are conserved host factors for CoV infection

Based on our initial gene enrichment results from the MERS-CoV and HCoV-229E screens, as well as a comparison of the respective results with previously published data^25–27^, we selected 21 hits for further experimental validation. Focusing on the highly pathogenic MERS-CoV screen, but also with an interest in examining common hits between both screens, we chose 17 genes that were significantly enriched in the MERS-CoV screen and 4 genes (TMEM41B, ELFN2, NOM1, and KRTAP13-4) that were significantly enriched in both MERS-CoV and HCoV-229E screens. For these 21 hits, stable CRISPR/Cas9 KO cell lines were generated for each gene and then challenged with either HCoV-229E or MERS-CoV. Specific KO of the MERS-CoV receptor DPP4 and the HCoV-229E receptor APN served as controls. MERS-CoV replication could be significantly reduced in all KO cell lines, except for WNT5A and APN, thus confirming our screen and validating our data analysis (Figures 3A and S4A). In contrast to MERS-CoV, HCoV-229E replication was significantly impaired upon deletion of APN as well as CDH7, MINAR1, TMEM41B, and FKBP8. Interestingly, KO of WNT5A significantly reduced HCoV-229E titers (Figures 3B and S4B). Importantly, TMEM41B, FKBP8, and MINAR1 knockout resulted in impaired titers for both MERS-CoV and HCoV-299E. Strikingly, SARS-CoV and SARS-CoV-2 also replicated to lower titers in respective KO cell lines expressing the specific entry receptor ACE2, confirming a conserved function in the CoV replication cycle for these three genes. (Figures 3C, 3D, and S4B). Western blot analysis confirmed stable knockout of both FKBP8 and TMEM41B (Figure 3E). Moreover, CRISPR/Cas9 mediated genome editing in MINAR1, FKBP8, and TMEM41B KO cell lines were confirmed via Sanger Sequencing (Figure S4E). To further validate the effect of the CRISPR/Cas9-mediated KO of all the three host factors, we expressed CRISPR resistant variants of these host factors and observed a rescue of virus titers for MERS-CoV, HCoV-229E, SARS-CoV, and SARS-CoV-2, thereby confirming the antiviral effect of TMEM41B, FKBP8, and MINAR1 KO (Figure 3F). To investigate the step of the viral replication cycle for which these factors are required, we employed a vesicular stomatitis virus (VSV) pseudo particle system bearing spike proteins from one of several different CoVs and encoding GFP as a reporter^28^. We found knockdown of TMEM41B, FKBP8, or MINAR1 did not alter VSV pseudoparticle entry mediated by spike proteins from HCoV-229E, MERS-CoV, SARS-CoV, or SARS-CoV-2 (Figure 4A). Collectively, these findings show that there is a conserved requirement for the host factors TMEM41B, FKBP8, and MINAR1 during CoV replication, but not during CoV entry.

**Figure 3.**
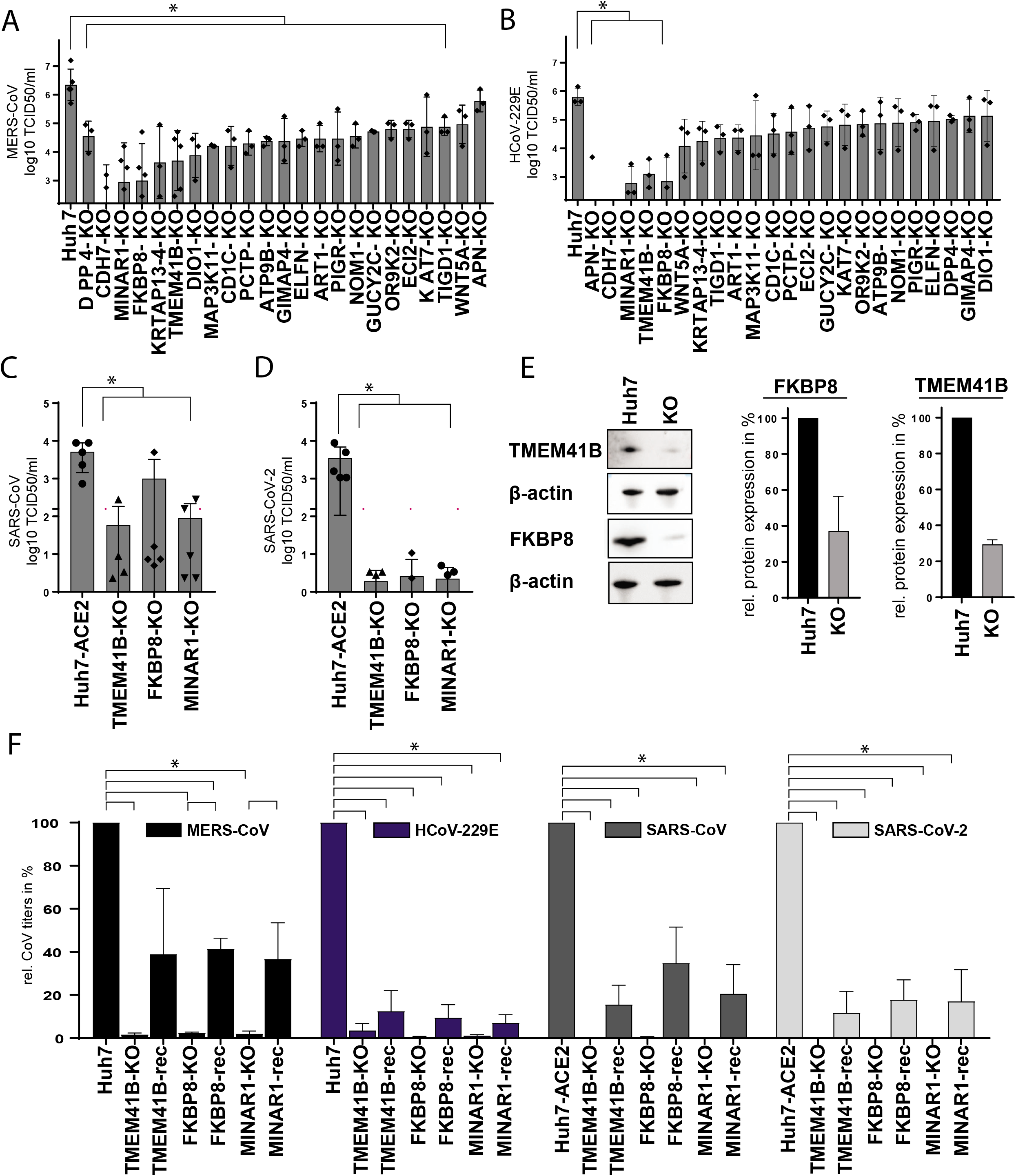
Top scoring host dependency factors are interactors of the autophagy pathway. MERS-CoV (A) and HCoV-229E (B) titers upon KO of top scoring HDFs are displayed in Log_10_ TCID_50_/ml. SARS-CoV (C) and SARS-CoV-2 (D) titers upon KO of TMEM41B, FKBP8 and MINAR1 are displayed in Log_10_ TCID_50_/ml. (E) Western Blot analysis of FKBP8-KO and TMEM41B-KO in Huh7 cells, including beta actin as loading control. MERS-CoV (F), HCoV-229E (G), SARS-CoV (H) and SARS-CoV-2 (I) titers upon reconstruction of TMEM41B, FKBP8 and MINAR1 in respective KO cell lines. Titers are shown relative to Huh7(-ACE2) control in %. Results are displayed as a mean of three with SD, represented by error bars. In A – D, statistical analysis was determined by ordinary one-way ANOVA, Dunnett’s multiple comparison test, using Nev 2020 version 9.0. In F, statistical significance was determined by two-tailed unpaired student t-test with Welch’s correction. Statistical calculations were performed in GraphPad Prism 8.3.1.

**Figure 4:**
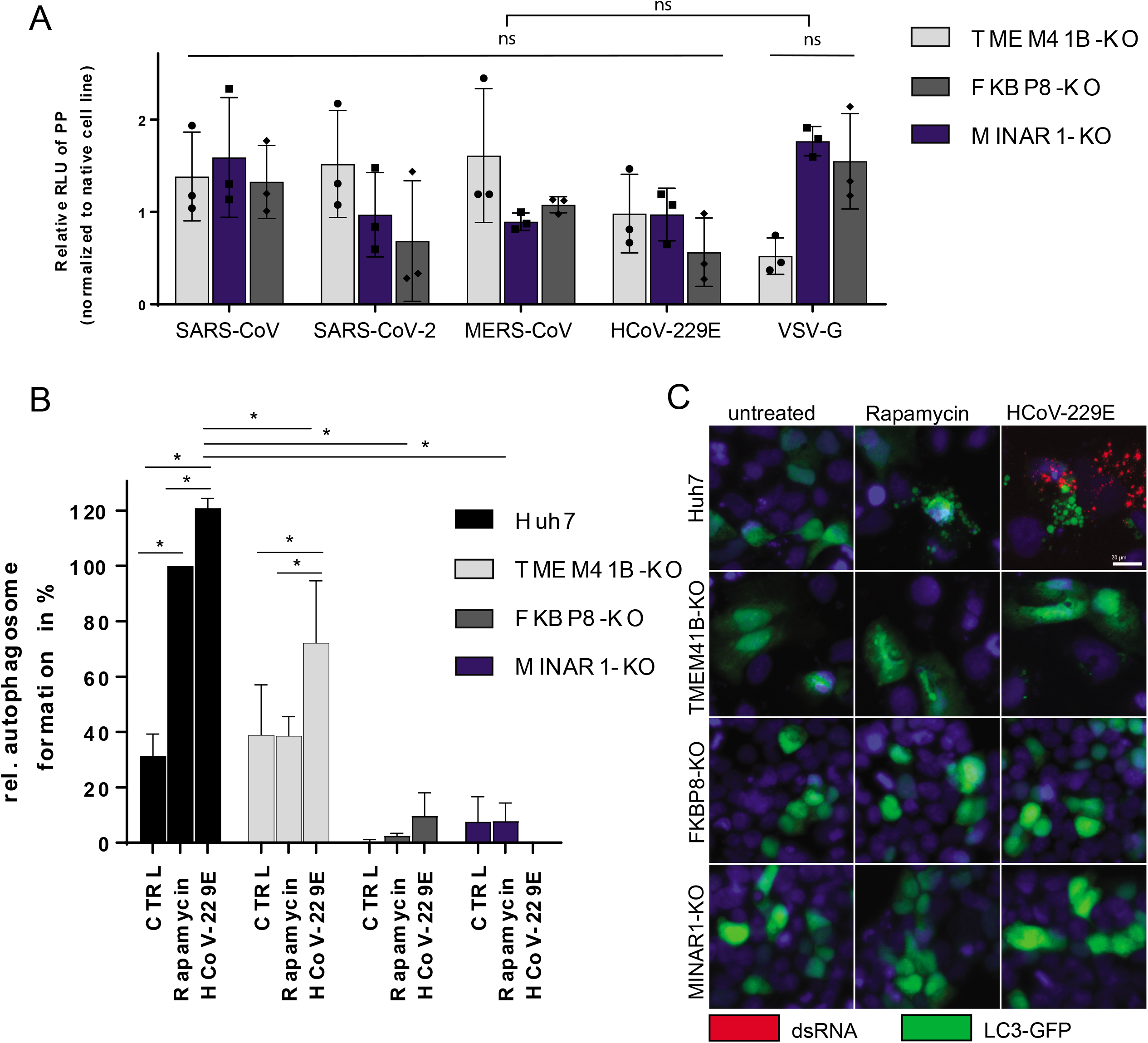
LC3-GFP translocation is impaired in TMEM41B, FBKP8 and MINAR1-KO cells. (A) VSV-G-based CoV-spike-mediated pseudo particle entry is shown in RLU in TMEM41B-KO, FKBP8-KO and MINAR1-KO cells normalized to entry in native cells. Results are displayed as a mean of three with SD, represented by error bars. One-way ANOVA statistical analysis was performed in Graph 8.3.1. (B) Quantification of (C) of LC3-GFP translocation shows relative autophagosome formation upon rapamycin treatment and HCoV-229E infection in native Huh7, as well as TMEM41B-KO, FKBP8-KO and MINAR1-KO cells. 5 images per condition in 3 independent experiments were acquired using an Evos FL Auto 2 imaging system with a 4x air objective, analyzed and quantified in Fiji. Statistical analysis was determined by two-tailed unpaired student t-test in GraphPad Prism 8.3.1. (C) Immunofluorescence staining of LC3-GFP expressing Huh7, TMEM41B-KO, FKBP8-KO and MINAR1-KO cells upon rapamycin treatment and HCoV-229E infection. LC3-GFP is depicted in green, dsRNA is shown in red and DAPI in blue, scale bar is 20 μm. Representative images of one out of four independent replications are shown. Images were acquired using an EVOS FL Auto 2 imaging system with a 20x air objective and processed using Fiji.

Despite having distinct cellular functions, TMEM41B, FKBP8, and MINAR1 are all involved in the cellular or mitochondrial autophagy pathways, albeit at different stages. As autophagy was also identified as one of the main conserved biological clusters in our GO analysis, we next chose to focus on these factors in the context of autophagy for further analysis. To confirm the association of TMEM41B, FKBP8, and MINAR1 with cellular autophagy, we induced autophagy in LC3-GFP transfected KO cells using Rapamycin and subsequently infected these cells with HCoV-229E. Under normal physiological conditions, the cytosolic protein LC3 translocates to autophagosomal membrane structures during early autophagy^29^. We thus analyzed the ability of LC3-GFP to translocate to such vesicles in TMEM41B, FKBP8, and MINAR1-KO cells infected with HCoV-229E and undergoing autophagy as described previously^29^ and analyzed our results using immunofluorescence (Figures 4B and 4C). In line with previous reports, we confirmed by visualizing LC3-GFP accumulation that Rapamycin treatment induced specific vesicle formation in native Huh7 cells, but not in TMEM41B-KO, FKBP8-KO, or MINAR1 KO cells, reasserting the necessity of these proteins for autophagosome formation. Similarly, LC3-GFP accumulated in Huh7 cells during HCoV-229E infection, but significantly less in TMEM41B-KO, FKBP8-KO, and MINAR1-KO cells (Figures 4B and 4C). Together these results show that KO of TMEM41B, FKBP8, and MINAR1 impairs membrane-remodeling during Rapamycin-induced autophagy and compromises LC3-GFP translocation during HCoV-229E infection.

### Inhibition of the immunophilin protein family with pre-existing drugs

TMEM41B, FKBP8, and MINAR1 have all been implicated as interactors of the autophagy pathway (Figure 5A). Moreover, FKBP8 is part of a large immunophilin family, known to bind the immunosuppressive agent Tacrolimus. Interestingly, in addition to FKBP8, several cyclophilins (additional members of the immunophilin family) were also significantly upregulated in the MERS-CoV and HCoV-229E CRISPR KO screens, including Peptidyl-prolyl isomerase (PPI) B, PPIC, PPID, PPIE, PPIF, PPIG and PPIH. Proteins of this family specifically bind Cyclosporin A, an immunosuppressant drug that is usually applied to suppress rejection after internal organ transplantation. Given the lack of specific treatment options for HCoVs, in particular during the ongoing SARS-CoV-2 pandemic, we tested Cyclosporin A, as well as Alisporivir, a non-immunosuppressant derivative of Cyclosporin A, currently used for treatment of HCV^30^. Importantly, both Tacrolimus and Cyclosporin A are known to bind and thereby inhibit calcineurin (PP3R1, MERS-CoV-specific HDF, Figure 5A) in their complexed form with the respective immunophilin^31^.

**Figure 5:**
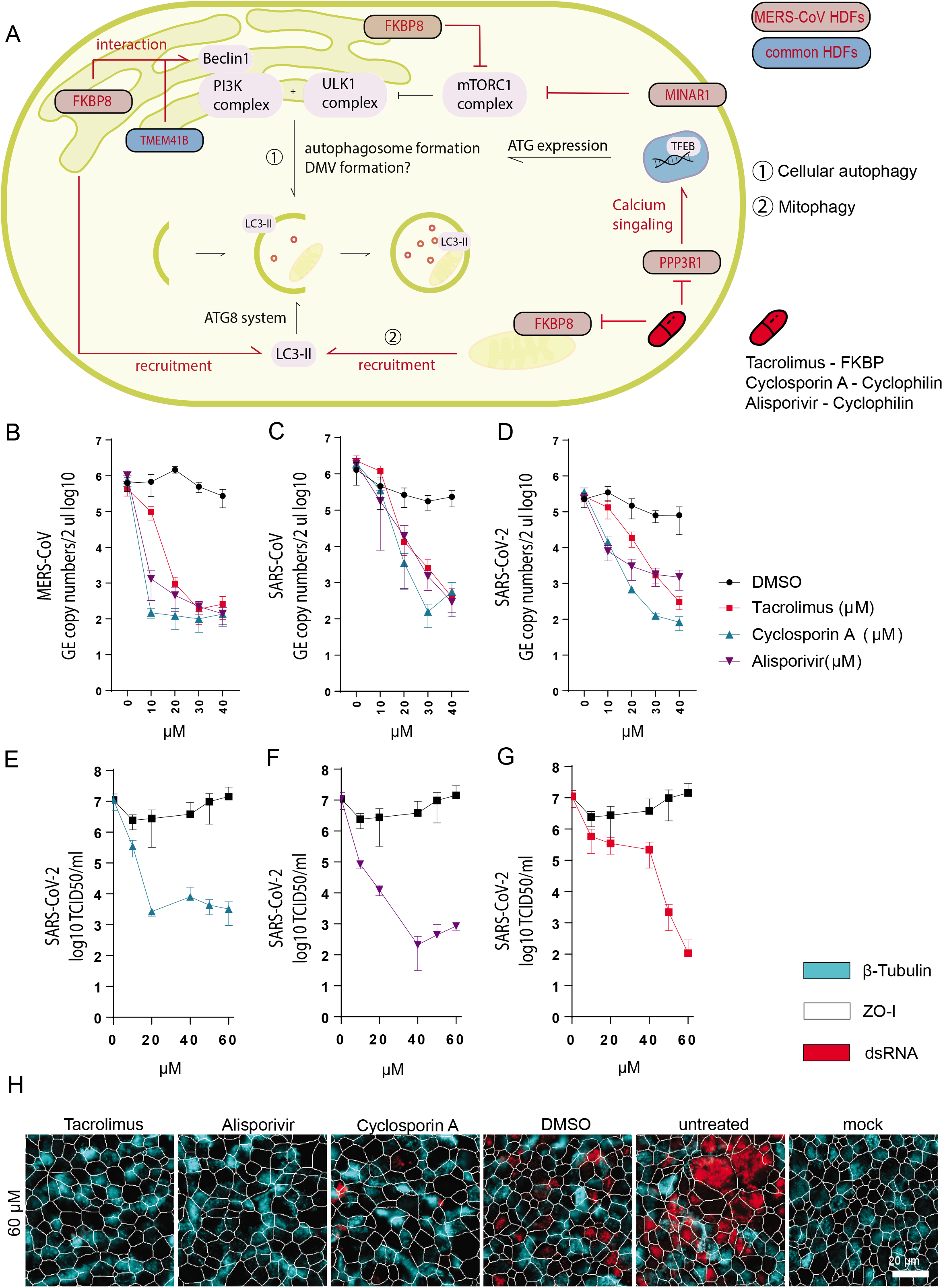
CoV HDFs are interactors of the autophagy pathway but do not depend on autophagy for replication. (A) Upon starvation, the mTORC1 complex is blocked and activation of the PI3K complex, as well as the ULK1 complex leads to the initiation of phagophore formation, as an initial step in the autophagy pathway. MERS-CoV and HCoV-229E top scoring CRISPR knockout screen hits FKBP8, MINAR1, TMEM41B and VMP1 are involved in this early pathway. Furthermore, the ATG8 system containing among others LC3, which is recruited by VPM1 or FBKP8 is necessary for targeting cellular cargo to the autophagosome. PPP3R1 is upregulated and initiates TFEB translocalization to the nucleus, where it catalyzes transcription of ATGs. MERS-CoV or conserved host dependency factors (HDFs) are indicated in respective colors. Inhibitor intervention in this pathway is shown in red. MERS-CoV (B), SARS-CoV (C) and SARS-CoV-2 (C) titers in TCID/ml upon treatment of Huh7 (MERS-CoV) and VeroE6 (SARS-CoV and SARS-CoV-2) cell lines with Tacrolimus, Cyclosporin A and Alisporivir at indicated concentrations. SARS-CoV-2 titers in TCID_50_/ml in primary human nasal epithelial cells at 48 hpi (E-G) and in presence of Cylosporin A, Alisporivir and Tacrolimus. (H) Immunofluorescence staining of SARS-CoV-2 infected human primary nasal epithelial cells, following DMSO, Tacrolimus, Alisporivir, Cyclosporin A and untreated treatment. dsRNA is shown in red, tight junctions (ZO-I) are shown in white and cilia (β-Tubulin) are shown in light blue. Images were acquired using an EVOS FL Auto 2 imaging system with a 40x air objective and processed using Fiji. dsRNA (red), cilia (β-Tubulin, light blue) and the outline of segmented cells (ZO-I, white) of representative images are shown. Scale bar: 20 μm.

Inhibitor treatment over the course of CoV infection resulted in a dose-dependent inhibition of MERS-CoV, SARS-CoV, as well as SARS-CoV-2 replication in cell lines 24 hours post infection. The most substantial reduction of genome equivalent copy numbers was up to 4 log upon Cyclosporin A treatment at concentrations starting at 10 μM for MERS-CoV (Figure 5B) and 30-40 μM for SARS-CoV (Figure 5C). Similar dose-dependence was observed for reduction of SARS-CoV-2 replication (Figure 5D). There are currently no drugs available against SARS-CoV-2. Therefore, we analyzed the effect of these compounds specifically against SARS-CoV-2 in a more biologically relevant cell culture system: primary well-differentiated human nasal epithelial cell cultures, which mimic the natural site of SARS-CoV-2 replication. Cyclosporin A inhibited SARS-CoV-2 replication at 48 hours post infection by around 4 log_10_ TCID_50_/ml at non-cytotoxic concentrations with a half maximal inhibitory concentration (IC_50_) of 7.9 μM (Figure 5E, S5E and S5J) and Alisporivir by approximately 4 log_10_ TCID_50_/ml at non-cytotoxic concentrations with an IC_50_ of 2.3 μM (Figure 5F, S5F and S5J). In contrast, the inhibition effect of Tacrolimus was accompanied by impaired cell viability in the nasal epithelial cell cultures (Figure 5G, S5G and S5J). Taken together, these findings suggest that these immunophilin interactors inhibit the function of certain HCoV HDFs, thereby impairing virus replication.

## Discussion

Identification of HDFs essential for HCoV infection offers great potential to reveal novel therapeutic targets and enhance our understanding of HCoV infection and pathogenesis (e.g., COVID-19). Here, we have performed two independent genome-wide CRISPR/Cas9 knockout screens in Huh7 cells with HCoV-229E and MERS-CoV to identify functionally important genes during HCoV infection. Using MERS-CoV as a representative emerging virus and HCoV-229E as a representative endemic virus, we identified multiple virus-specific and conserved HDFs, including several that are required for the replication of the novel pandemic CoV SARS-CoV-2. GO enrichment analysis revealed that the conserved HDFs were involved in diverse biological processes that clustered into seven major categories. Interestingly, we found that MERS-CoV and HCoV-229E seemed to exploit different components of the same biological processes, as the majority of genes involved in each biological cluster were virus specific, but the overall biological processes were similar. This may be due to evolutionary differences between the viruses, as MERS-CoV is part of the betacoronavirus genus whereas HCoV-229E is a member of alphacoronavirus genus. Furthermore, many commonly enriched genes were involved in Golgi vesicle transport, or more specifically in vesicle coating and budding from membranes, as well as regulation of endocytosis and exocytosis, which are known to be associated with virus entry and exit^32^. Moreover, Golgi vesicle markers have been found in close proximity to CoV replication compartments, suggesting another potential function for genes in this cluster during CoV replication, e.g. membrane re-organization for membranous replication compartments^33^. A second prominent category was the immune system cluster, which may be associated with direct exploitation of immunological host responses against CoVs and thus offer potential intervention strategies. These strategies may also have antiviral efficacy and work to lower dysfunctional immune responses, which is a known driver of disease progression and severe lung pathology^34^. Another major category containing enriched genes in both HCoV screens was dephosphorylation. Genes involved in phosphorylation and kinase activities were strongly enriched in our screens, suggesting that these processes are required for HCoV replication and that other CoVs also exploit the host’s phosphorylation machinery for their benefit. Importantly, recent work observed striking changes in phosphorylation on host and viral proteins during SARS-CoV-2 infection, including many changes related to dephosphorylation and altered kinase activity^35,36^. For example, the JNK signaling cascade, but also the regulation of tau-protein kinase activity, were highly enriched in our MERS-CoV screen. JNKs belong to the mitogen-activated protein kinase (MAPK) family and SARS-CoV-2 infection was recently shown to promote p38 MAPK signaling activity^35^. Of note, the FKBP8 gene clustered into the dephosphorylation category and the MINAR1 gene was included in regulation of tau-protein kinase activity, suggesting that these two genes may influence CoV replication via other biological processes in addition to autophagy. Along this line, therapeutical intervention targeting AP2M1 (part of the clathrin-dependent endocytic pathway) phosphorylation using a kinase inhibitor resulted in reduced SARS-CoV, MERS-CoV and SARS-CoV-2 infection, exemplifying the antiviral potential of targeting specific phosphorylation sites during viral infection^37^. Finally, our analysis also found that genes involved in catabolic and homeostatic processes were significantly enriched in both CoV screens. Interestingly, a similar cluster linked to cholesterol metabolism was identified in previous studies, including SARS-CoV-2, HCoV-229E, and HCoV-OC43 genome-wide CRISPR/Cas9-mediated KO screens and SARS-CoV-2 interactome studies^21,38^ and has been linked to CoV entry and membrane fusion^39^.

For our downstream experimental analysis, we focused on the autophagy cluster. Autophagy is a cellular stress response to e.g. starvation or infection by pathogens for the recycling of proteins and cell organelles to maintain cellular homeostasis^40^. The processes comprises a very wide-ranging family of trafficking pathways required for the transportation of cytoplasmic material to the lysosome for destruction. The ER localized TMEM41B was recently identified as a gene required for early autophagosome formation and lipid mobilization in three independent genome-wide CRISPR knockout screens, which also observed that TMEM41B and the well-characterized early-stage autophagy protein VMP1 (top scoring HDF in HCoV-229E screen) implement related functions^25,26,27^. Furthermore, interaction of TMEM41B with Beclin1 (PI3K complex) underscores the importance of this protein in the induction of autophagy^41^. Interestingly, the FK506-binding protein 8 (gene: FKBP8, protein: FKBP38), a member of the immunophilin protein family is located in the outer mitochondrial membrane and plays a key role in mitophagy by inhibiting the mTORC1 complex during nutrient deprivation^42^. Moreover, FKBP8 targets Beclin-1 to ER-mitochondria membranes during mitophagy and recruits LC3A to damaged mitochondria, thereby actively inducing the removal of excess mitochondria by autophagy^43^. FKBP8 itself avoids degradation by escaping from mitochondria and is translocated to the ER^44^. MINAR1 (also known as Ubtor or KIAA1024) was the third MERS-CoV HDF with a possible indirect involvement in autophagy regulation. The otherwise very rudimentary characterized protein plays a role in regulating cell growth and mTOR signaling, as MINAR1 depletion resulted in higher mTOR activity^45^ (Figure 5A). In addition, the phosphatase PPP3R1, commonly referred to as calcineurin, is upregulated during cell starvation and controls the activity of the TFEB transcriptional regulator of lysosomal biogenesis and autophagy^46^. Importantly, the interaction between autophagy components and CoVs but also other positive-stranded RNA viruses during viral replication has been under discussion for a long time, as parts of the autophagy process show similarities to the process of DMV formation^47,33,48^. CoVs rely on the formation of replication complexes at DMVs, the presumed site of viral genome replication and transcription. Due to a lack of conventional endoplasmic reticulum (ER) or Golgi protein markers the exact origin of DMVs remains unclear and studies investigating the possible involvement of the early autophagy machinery in the conversion of host membranes into DMVs reached conflicting conclusions^49,50^. Another possibility is that single components of the autophagic machinery may be hijacked by CoVs independently of their activity in autophagic processing. The non-lipidated autophagy marker LC3 has been observed to localize to DMVs and the downregulation of LC3, but not inactivation of host cell autophagy, protects cells from CoV infection^51,52,47,53^. We show that TMEM41B, MINAR1 and FKBP8 are involved in regulating vesicle formation during autophagy as LC3-GFP did not relocate to characteristic foci indicative of autophagosomes following chemical induction of autophagy and that KO of each gene distinctly impairs HCoV replication, but the mechanistic connection of both processes remains elusive. Further roles of the three identified host factors have been suggested. Both TMEM41B and FKBP8 are thought to interact with Beclin-1, which is a core subunit of the PI3K complex that drives autophagy^41^,^54^. Captivatingly, inhibition of SKP2, another Beclin-1 interactor, reduced MERS-CoV infection.^55^ Recent work suggested a putative autophagy-independent role for TMEM41B as a pan-coronavirus and flavivirus replication factor, which is recruited to flavivirus RNA replication complexes to facilitate membrane curvature and create a protected environment for viral genome replication^56,20^. Furthermore, MINAR1 serves as a regulator of mTOR signaling, which regulates numerous cellular processes including the cap-dependent mRNA translation and synthesis machinery required during viral replication. These observations add further potential layers of modulation by TMEM41B, FKBP8 and MINAR1 during CoV replication.

Independently of the exact underlying mechanism, our results suggest that the HDFs FKBP8, TMEM41B, and MINAR1 herein represent potential targets for host-directed therapeutics. Its immunomodulating component, make FKBP8 a very interesting HDF for CoV replication. FKBP8 is part of the immunophilin family of FK506-binding proteins, which share the ability to act as a receptor for the immunosuppressive drug FK506 (Tacrolimus), usually used to lower the risk of transplant rejection after allogenic transplantation^57^. On a different note, knockdown of FKBP8 promotes the activation of IFN-beta and the antiviral response during Sendai virus infection in HEK293T cells, suggesting a possible immunomodulatory component for its role in CoV infection^58^. In addition to FKBP8, cyclophilins were upregulated in both HCoV screens. Cyclophilins express PPI activity, which catalyzes the isomerization of peptide bonds in proline residues from *trans* to *cis*, thereby facilitating protein folding. Proteins of this family specifically bind Cyclosporin A, an immunosuppressant drug that is usually applied to suppress rejection after internal organ transplantation. Moreover, immunophilins and cyclophilin have been in the focus of several CoV studies showing impaired HCoV-229E, HCoV-NL63, as well as SARS-CoV and MERS-CoV replication upon immunophilin and cyclophilin inhibitor treatment^59,60,61,62,63,64^. Given the lack of specific treatment options during the ongoing SARS-CoV-2 pandemic, we tested Tacrolimus, Cyclosporin A, as well as Alisporivir, a non-immunosuppressant derivative of Cyclosporin A and showed that antiviral intervention using these clinically approved immunosuppressive drugs inhibited the replication of the highly pathogenic CoVs MERS-CoV, SARS-CoV, and SARS-CoV-2 in a dose-dependent manner. While Huh7 and VeroE6 cells are valuable model cell lines for highly pathogenic CoVs, they likely do not capture important aspects of infection compared to primary human airway epithelial cells nor fully recapitulate the complex cellular milieu present in human patients. To address these limitations, we also tested these drugs on primary human nasal epithelial cell cultures and found that both Alisporivir and Cyclosporin A potently inhibit SARS-CoV-2 replication at concentrations known to be achievable and efficacious in patients. Together these findings depict a promising path towards the repurposing of Cyclosporin A and Alisporivir as COVID-19 treatment options. Infection with highly pathogenic CoVs is frequently accompanied by inflammatory immunopathogenesis, including the virus-induced destruction of lung tissue and subsequent triggering of a host immune response. Importantly, in certain cases a dysregulated immune response is associated with severe lung pathology and systemic pathogenesis^34^. The latter highlights the need for dual-acting antiviral drugs that also target inflammation and/or cell death. Of interest, Alisporivir also blocks mitochondrial cyclophilin-D, a key regulator of mitochondrial permeability transition pore (mPTP) opening, which is a mechanism involved in triggering cell death. Hence, besides its antiviral properties, it is possible that Alisporivir also reduces CoV-induced lung tissue damage^65^. Trials using either Cyclosporin in patients with moderate COVID-19 (ClinicalTrials.gov Identifier: NCT04412785 and NCT04540926) or Alisporivir (ClinicalTrials.gov Identifier: NCT04608214) for the treatment of hospitalized COVID19 patients have been registered.

The identification of MINAR1, TMEM41 and FKBP8 as conserved HCoVs HDFs in our MERS-CoV and HCoV-229E screens extend the knowledge on HCoVs. Furthermore, the involvement of FKBP8 and other members of the cyclophilin family in the HCoV replication provide information on how Tacrolimus, Cyclosporin A and Alisporivir are able reduce CoV replication by interfering with essential HCoV HDFs. We confirm the potential of all three inhibitors as treatment against HCoV infections, and additionally observed similar reduction in SARS-CoV-2 replication. Altogether our findings highlight the potential of genome-wide CRISPR/Cas9 knockout screens to identify novel HDFs essential for HCoV infection, which can in turn be used in combination with clinically available drugs to identify and evaluate host-directed therapies against existing and future pandemic CoVs.

## Author Contributions

AK: experimental setup, data collection, data analysis; writing; JK: experimental setup, data analysis, writing; YB: writing; JP: data collection; PV. data analysis; DT: data analysis. NE: data collection; ES: experimental setup; RD: experimental setup, data analysis; GZ: experimental setup, data collection, SP: experimental setup, data collection, data analysis; writing; VT: experimental setup, writing.

## Acknowledgements

This study was supported by Swiss National Science Foundation (SNF; grants 165076 and 173085), the European Commission (Marie Skłodowska-Curie Horizon 2020 project “COV RESTRICT” grant agreement No. 748627), the Federal Ministry of Education and Research (BMBF; grant RAPID, #01KI1723A) We would like to thank the Next Generation Sequencing Platform of the University of Bern. We would like to thank Biosafety at the Institute for Virology and Immunology (IVI) in Mittelhäusern. We would like to thank for the support of all members of the IVI Bern and the Ruhr-University in Bochum. Finally, we are grateful to Christian Drosten and Marcel Müller for the virus isolates.

## Main tables

**Table 1: MAGeCK results for MERS-CoV and HCoV-229E screens**

**Table 2: GO term analysis results**

## Supplemental figure titles and legends

**Figure S1: Quality control metrics and enriched gene identification for MERS-CoV and HCoV-229E genome-wide CRISPR screens.** (A) Area under the curve (AUC) analysis of MERS-CoV and HCoV-229E CRISPR screens evaluating sgRNA library representation in surviving Huh7 cells from uninfected (Mock) and MERS-CoV (left two panels) or HCoV-229E (right two panels) infected samples. For each CRISPR screen, sgRNA abundance was calculated based on average sgRNA abundance over 3 independent biological replicates. (B) Correlation matrix depicting the Pearson correlation for guide-level normalized read counts among biological replicates and samples from both screens. R1, R2, and R3 represent the biological replicates 1, 2, and 3, respectively. Clustering was performed in pheatmap using correlation as a distance metric (C) Robust Rank Aggregation (RRA) p-value distribution of all genes in the GeCKOv2 library for both MERS-CoV (left) and HCoV-229E (right) CRISPR screens. Genes that met the criteria for significance (RRA p-value ≤ 0.05 and FC ≥ 2) are highlighted in red. (D) Venn diagram illustrating the overlap between significantly enriched genes from both CRISPR screens that were identified via two different RRA-based analysis methods (alpha median and second best). A total of 19 genes were identified by both methods in both MERS-CoV and HCoV-229E CRISPR screens.

**Figure S2**: (A) Representative GO terms identified using full list of enriched GO terms for MERS-CoV and HCoV-229E screens (Table 2). Representative terms found in both screens are shown in the top panel, whereas virus-specific terms are shown in the bottom panel. BP, CC, and MF represent different GO term categories. (B) Specific GO terms enriched in both CoV screens (individual GO terms, not representative GO terms).

**Figure S3:** Cnet plots for GO BP terms found in each individual biological cluster shown in Figure 2A (S3A Golgi Vesicle Transport, S3B Autophagy, S3C Catabolic Processes, S3D Dephosphorylation, S3E Immunity, S3F Developmental Processes, S3G Homeostatic Processes). Plots include both GO terms that contain one or more of the 19 common significantly enriched genes found in both CoV screens (as in Figure 2A and 2B) as well as representative GO terms found in both screens that do not contain these genes. Each plot shows the relationship among individual GO terms and genes found in each biological cluster. Larger nodes represent individual GO terms and smaller nodes represent individual gene. Nodes that are functionally related cluster together into a larger network. Node size reflects the number of significantly enriched genes in the node and color indicates the CoV screen for which the node was significant.

**Figure S4: CRISPR-mediated KO of top scoring host dependency factors impairs CoV replication.** (A) Immunofluorescence staining of MERS-CoV infected of Huh7 cells containing KO of top scoring HDFs. dsRNA is shown in green, DAPI is shown in blue. (B) Immunofluorescence staining of HCoV-229E, SARS-CoV and SARS-CoV-2 infected Huh7 cells with TMEM41B, FKBP8 and MINAR1-KO, as well as a stable ACE2 expression. dsRNA is shown in green, DAPI is shown in blue, ACE2 is shown in red. Scale bar is 50 μm. All images were acquired using an Evos Auto FL2 and processed in Fiji. (D) Relative cytotoxicity of TMEM41-KO, FKBP8-KO and MINAR1-KO is depicted in %. Two tailed unpaired student t-test was used to determine significance in GraphPad Prism 8.3.1. (E) Sanger sequencing of FKBP8-KO, MINAR1-KO and TMEM41B-KO verifies Cas9-mediated double strand break in multiple alleles of the KO cells. PAM sequence is indicated in red, binding site of sgRNA is indicated in blue.

**Figure S5: Cyclosporin A, Alisporivir and Tacrolimus inhibit CoV infection in a dose-dependent manner in cell lines and primary human nasal epithelial cells at non-cytotoxic concentrations.** Immunofluorescence staining of SARS-CoV (A), as well as SARS-CoV-2 (B) infected VeroE6 cells and MERS-CoV (C) infected Huh7 cells following Cyclosporin A, Alisporivir, Tacrolimus treatment at 10 μM to 40 μM and Bafilomycin treatment at 10 nM – 40 nM, as well as well as DMSO CTRL as respective volumes 24 hrs post infection/inhibitor treatment. dsRNA is shown in green, DAPI is shown in blue. Scale bar is 50 μm. (D) Immunofluorescence staining of SARS-CoV-2 infected and Cyclosporin A, Alisporivir and Tacrolimus, as well as DMSO treated primary human nasal epithelial cells at 10 μM to 60 μM 48 hpi/post inhibitor treatment. dsRNA (red), cilia (β-Tubulin, light blue) and the outline of segmented cells (ZO-I, white) are shown. Scale bar is 20 μm. All images were acquired using an EVOS FL Auto 2 imaging system with a 10x air objective (A, B, C) and a 40x air objective (D) and processed using Fiji. (E-G) Inhibitor treated primary nasal epithelial cell cultures displayed as inhibitor versus normalized response. IC_50_ value is marked with dotted line and indicated on y axis. Calculations were performed in GraphPad Prism 8.3.1. Cyclosporin A, Alisporivir andTacrolimustreatment-mediated cytotoxicity in Huh7 cells (H) and VeroE6 cells (I) shown relative to dead cell control. (J) Relative nasal epithelial cell culture viability upon treatment of (50 μM and) 60 μM Cyclosporin A, Alisporivir and Tacrolimus normalized to DMSO.

## Methods

### Lead Contact

Further information and request for resources and reagents should be directed to and will be fulfilled by Volker Thiel. Unique reagents generated in this study will be made available on request.

### Materials Availability

Unique reagents generated in this study will be made available on request. This applies to pCaggs-MINAR1mut, pCaggs-FKBP8mut, with silent mutations in Cas9 binding PAM region, as well as pCaggs-TMEM41Bmut. Payment/MTA may be required.

### Data and Code Availability

Sequencing data from CRISPR/Cas9 knockout screens will be made available in a public repository upon publication.

### Experimental Model and Subject Details

#### Cell Lines

Human hepatoma (Huh7) cell line (kindly provided by Volker Lohmann) and African green monkey kidney (VeroE6) cell line (kindly provided by Doreen Muth, Marcel Müller and Christian Drosten, Charité, Berlin, Germany) and 293LTV cells (purchased from Cell Biolabs Inc.) were propagated in Dulbecco’s modified Eagle Medium (DMEM), supplemented with 10% heat inactivated fetal bovine serum, 1% nonessential amino acids, 100 μg/mL of streptomycin and 100 IU/mL of penicillin, and 15 mMol of HEPES. Cells were maintained at 37°C in a humidified incubator with 5% CO2. Profiling of cell lines was performed using highly-polymorphic short tandem repeat loci (STRs) and amplification using PowerPlex 16 HS System (Promega), followed by fragment analysison an ABI3730xl (Life Technologies) and analysis with GeneMarker HID software (Softgenetics) by Mircosynth. Huh7 cell line was confirmed to be of human origin without contamination, matching the reference DNA of the cell line Huh7 (Microsynth reference, Mic_ 152021) with 96.7 % and the DNA profile of Huh7 (Cellosaurus, RRID:CVCL_0336) with 90 %. 293 LTV cell line was confirmed to be of human origin without contamination, matching the reference DNA of the cell line HEK293T (ATCC^®^ CRL-3216™) with 93.8 % and the DNA profile of HEK293 with 86.7 % (Cellosaurus, RRID:CVCL_0045). Matching at ≥ 80 % of alleles across eight reference loci are said to be related. VeroE6 cell line was identified to be 100% identical with Chlorocebus sabaeus, upon amplification and blast of mitochondrial cytochrome b gene according to DM Irwin *et al*.^69^, using primers:

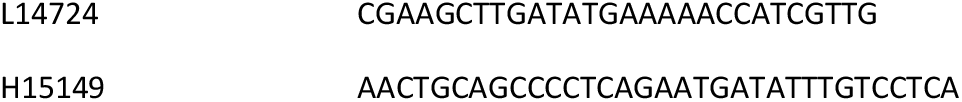

#### Primary Cell Culture

Primary human nasal epithelium cell cultures: MucilAir™ were purchased from epithelix. Cultures are reconstituted using human primary cells from healthy nasal region from 14 donors and cultured at an air-liquid interface in ready-to-use MucilAir™ Culture Medium purchased from epithelix is serum free, contains phenol red and is supplemented with penicillin/streptomycin. The apical side was washed with HBBS prior to infection. The anonymity of the donors prevents from the determination of the cells’ sex.

### Method Details

#### Genome-wide CRISPR/Cas9-mediated Knockout Screens

The vector lentiviral human GeCKOv2 library A^70^, containing 3 sgRNAs per gene, was transfected into 293 LTV cells for lentivirus production using Lipofectamine 2000 (Thermo Fisher Scientific). The supernatant was collected 48 hours post transfection and clarified by centrifugation (3500 *rcf*, 15 min). Huh7 cells were subsequently transduced with GeCKO lentiviruses at a MOI of 0.3 and selected for with puromycin at a concentration of 0.25 μg/ml for 7 days. To ensure sufficient sgRNA coverage, 60 Mio selected Huh7 cells were infected with either HCoV-229E (33°C, MOI 0.1) or MERS-CoV (37°C, MOI 0.05) and then incubated until the non-transduced control cells died. Non-transduced Huh7 cells were infected with respective viruses to control for complete cytopathic effect. Both screens were performed in 3 independent biological replicates. Surviving cells were harvested approximately two weeks post infection and genomic DNA was isolated using the Macherey Nagel NucleoSpin Tissue Kit according to the manufacturer’s instructions. All sgRNAs were amplified from genomic DNA using a two-step PCR protocol, enabling multiplexing and the addition of specific barcodes for Illumina sequencing on a NovaSeq using 60 Mio reads and paired end reads 150. Illumina Adapter Primers^71^:

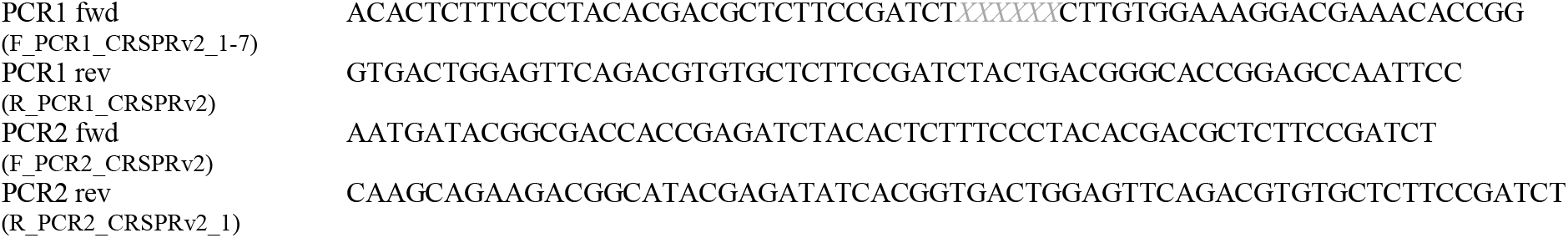

PCR products were then purified using Macherey Nagel PCR Clean Up pooled, and sequenced on the Illumina NovaSeq 6000 at the Next Generation Sequencing (NGS) facility at the University of Bern. The input library was also sequenced using the Illumina NGS platform to ensure full representation of sgRNAs in the GeCKO library.

#### Computational Analysis of Genome-wide CRISPR/Cas9-mediated Knockout Screens

Demultiplexed FASTQ files were trimmed and aligned to the reference sequences in the sgRNA library file. sgRNA abundance was quantified using the ‘count’ command from the MAGeCK pipeline and counts were compared between uninfected and infected samples to determine positive enrichment scores for each gene. MAGeCK testing was performed using paired analysis with the ‘alpha mean’ and ‘second best’ methods. Genes with a Robust Rank Aggregation (RRA) p-value of ≤ 0.05 and a log fold change (LFC) of ≥ 2 were considered significantly enriched. For both the MERS-CoV and HCoV-229E screens, data from three independent biological replicates was used as the input for data analysis. The gene ontology (GO) enrichment was performed on significantly enriched genes from each CoV screen using the ‘compareCluster’ function in clusterProfiler with the ‘fun’ option set to “enrichGO” and a formula of “Entrez ~ Screen”. To reduce GO term redundancy and identify a representative GO term for groups of similar terms, the rrvgo package was used in R with the similarity threshold set to 0.75. Finally, the plot in Figure 2A was created using the ‘emapplot_cluster’ function in the enrichplot package with a filtered version of the compareCluster enrichment result (filtered to include representative GO terms found in both CoV screens that contained one or more of the 19 common significantly enriched genes). All heatmaps were generated using the pheatmap package in R with clustering distance set to “Euclidean” and using the complete linkage clustering method. Volcano plots and venn diagrams were created using the EnhancedVolcano and VennDiagram packages, respectively.

### Characterization and Analysis of Top Scoring Host Dependency Factors

#### ACE2 Expression, FKBP8, TMEM41B and MINAR1 KO in Huh7 Cells

pSCRPSY-Tag-RFP-ACE2 (kindly provided by John Schoggins) was used for lentivirus production as described above and Huh7 cells were transduced and selected for using 0.5 ug/ml Blasticidin. ACE2 expression was confirmed via RFP expression. sgRNAs with highest scores in CRISPR-KO screen were ordered as forward and reverse oligos for creation of stable knock-out cell lines.

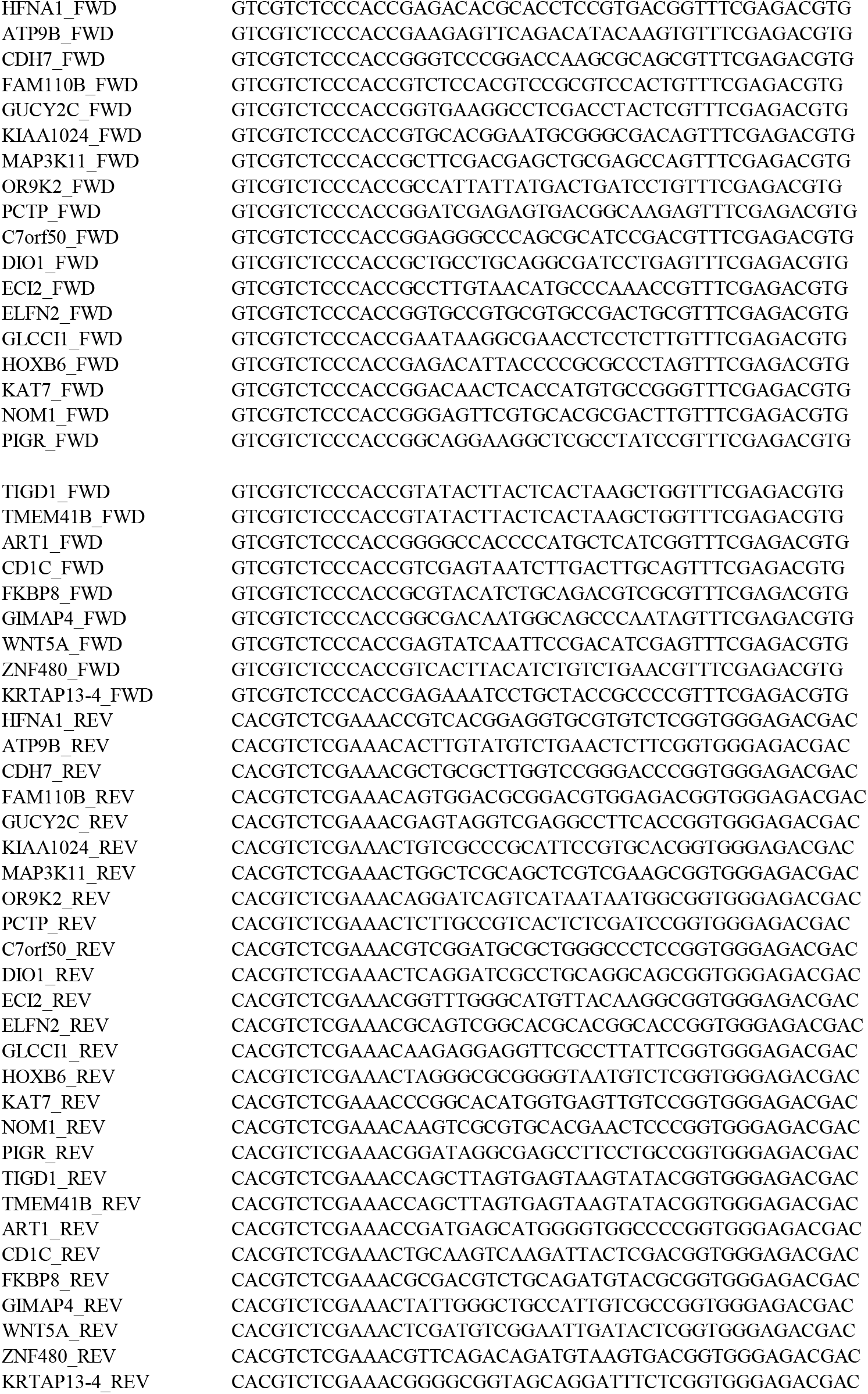

Oligonucleotides were denatured for 5 min at 99°C in TE buffer and then slowly adapted to room temperature and assembled with pLentiCRISPRv2 vector using Golden Gate cloning.

Plasmids were transformed in Stellar cells (Takara) and prepped for sanger sequencing and lentivirus production. ACE2-expressing Huh7 cells were transduced with pLentiCRISPRv2 containing sgRNAs for top scoring hits and selected with 0.25 ug/ml puromycin. Bulk knock-out of FKBP8, TMEM41B and MINAR1 was verified using Sanger Sequencing and Western Blot.

#### Western Blot

500.000 cells were lysed in M-PER Mammalian Protein Extraction Reagent (Thermo Scientific 78501) containing 1x protease inhibitor (cOmplete Tablets, Mini EDTA-free, EASYpack, Roche, 04693159001), mixing at 600 rpm for 10 min at RT in a ThermoMixer. Lysed cells were denatured with SDS at 95°C for 5 min and separated on an 10% SDS PAGE (SurePAGE Bis-Tris, 10×8, GenScript, M00666) at 200V for 30 min. eBlot L1 –Fast Wet Protein Transfer System (GenScript) was used for blotting and proteins were stained using the following antibodies: (Sigma-Aldrich AV46863), TMEM41B (Sigma-Aldrich HPA014946), MINAR1 (Sigma-Aldrich HPA011545), β-Actin-HRP (Sigma, A3854), as well as donkey anti rabbit-HRP (JacksonImmunoResearch, 711-035-152). Proteins were visualized using WesternBright ECL HRP substrate (Advansta, K-12045-D20) and the Fusion FX (Vilber) imaging system.

#### VSV Pseudotype Particles Bearing CoV Spike Proteins

Approximately 6*10^5^ 293LTV cells were seeded into a six-well plate and transfected with expression plasmids encoding either VSV-G surface protein (positive control, VSV-G; GenBank accession number NC_001560), HCoV-229E spike (pCAGGS-229E S; GenBank accession number X16816), MERS-CoV spike (pCAGGS-MERS S; GenBank accession number JX869059, with a silent point mutation (C4035A, removing internal XhoI)), SARS-CoV spike (pCAGGS-SARS S; GenBank accession number: AY291315.1, with two silent mutations (T2568G and T3327C)) or SARS-CoV-2 spike (generated as described^28^) using the transfection reagent Lipofectamine 2000 as described previously ^28^,. At 20 hours post transfection, cells were infected with VSV-G-trans-complemented VSV*ΔG(FLuc) (MOI = 5) at 37 °C. After inoculating the cells with virus for 30 min, they were washed with PBS and incubated for 24 hours with DMEM medium containing a monoclonal neutralizing monoclonal antibody directed to the VSV-G protein (antibody I1, ATCC, 1:100). The cell culture supernatant was harvested and cleared by centrifugation (3,000g for 10 min) and used to inoculate Huh7 native and knockout cell lines for 24 hours, prior to measurement of luciferase using Bright-Glo Luciferase Assay System (Promega, E2620) and using a plate luminometer (EnSpire 2300 Multilabel reader; Perkin Elmer).

#### Viruses

HCoV-229E^66^ was propagated on Huh7 cells. MERS-CoV strain EMC^67^ was propagated in VeroB4 cells. SARS-CoV strain Frankfurt-1^68^ and SARS-CoV-2 (SARS-CoV-2/München-1.1/2020/929, kindly provided by Daniela Niemeyer, Marcel Müller and Christian Drosten) were propagated on VeroE6 cells.

#### Virus Infection

Huh7 cells were plated to 15.000 cells and VeroE6 cells were plated to 20.000 per 96 well 24 hours prior to infection. Cells were infected with HCoV-229E (33°C), MERS-CoV (37°C), SARS-CoV (37°C) and SARS-CoV-2 (37°C) at an MOI of 0.01 (MOI 0.1 for HCoV-229E) for 2 hours. The virus inoculum was removed and cells were washed 3 times with PBS. Primary human nasal epithelial cell cultures were infected with SARS-CoV-2 at an MOI of 0.1 at 37°C for 1 hour from the apical side. Inoculum was removed and cell 3 times with HBBS. In case of inhibitor treatment, Tacrolimus, Cyclosporin A or Alisporivir were added to the cell supernatant/basolateral medium directly after the removal of the inoculum and the washing of the cells at following concentrations: 0 uM, 10 uM, 20 uM, 30 uM, 40 uM, 50 uM, 60 uM. DMSO solvent control was added at respective volumes. The inhibitor was not removed during the course of infection. At 24-48 hours post infection the cells/supernatant were/was harvested and analyzed using titration, immunofluorescence staining or quantitative RT-PCR.

#### Virus Titration

In order to determine the 50% tissue culture infectious dose (TCID_50_) per milliliter (apical) supernatant was serially diluted at indicated hours post infection, Huh7 (MERS-CoV, HCoV-229E) VeroE6 cells (SARS-CoV(−2)) were inoculated with serial dilution and TCID_50_ per milliliter was visualized using Crystal Violet and calculated by the Spearman-Kärber algorithm after 72 hrs −120 hrs as described^72^.

#### Quantitative RT-PCR

Virus replication was analyzed via qRT PCR, viral RNA was isolated from the supernatant at indicated hours post infection using the NucleoMag Vet Kit (Macherey Nagel) and a Kingfisher Flex Purification System (Thermo Fisher Scientific, Darmstadt, Germany) according to manufacturer’s guidelines. Extracted RNA was amplified using TagMan™ Fast Virus 1-Step

Master Mix (Thermo Fisher Scientific). Following primers were used for detection of MERS-CoV^73^:

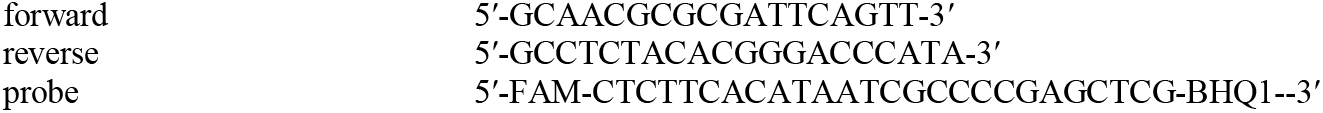

SARS-CoV and SARS-CoV-2:

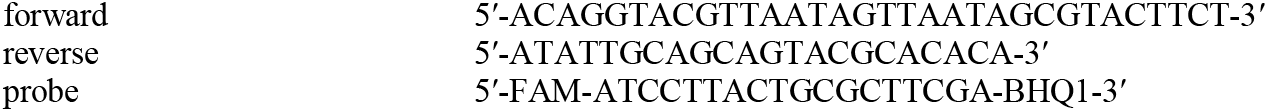

targeting the Envelope gene of SARS-CoV-2 (MN908947.3) The primers were adapted from Corman and colleagues^74^. A serial dilution of *in vitro* transcribed MERS-CoV RNA (kindly provided by Marcel Müller and Christian Drosten)^73^ and RdRp-E-N RNA mixture derived from a SARS-CoV-2 synthetic construct (MT108784) was included to determine the genome copy number^75^. Five *in vitro* transcribed (IVT) RNA preparations were produced from five different DNA fragments to cover the regions used for real-time RT-qPCR methods for the detection of SARS-CoV-2 and SARS-CoV viral RNA. Measurements and analysis were performed with the Applied Biosystems™ 7500 Fast Dx Real-Time PCR Systems and associated software (Applied Biosystems, Foster City, CA, USA).

#### Immunofluorescence Staining

For immunofluorescence staining cells were fixated with 4% formalin. Fixated cells were permeabilized in PBS supplemented with 50 mM NH4Cl, 0.1% (w/v) Saponin and 2% (w/v) Bovine Serum Albumin and stained with a mouse monoclonal antibody against dsRNA (SCICONS, clone J2). Alexa-Fluor 488-labeled donkey-anti mouse IgG (H+L) (JacksonImmuno, 715-545-150) was used as a secondary antibody. Alexa-Fluor^®^ 647-labelled rabbit anti-beta-tubulin IV (Cell Signalling Technology, 9F3) and Alexa-Fluor^®^ 594-labelled mouse anti ZOI-1 (Thermo Fisher Scientific, 1A12) were used to visualize cilia and tight junctions in nasal epithelial cell cultures. Cells were counterstained using 4’,6-diamidino-2-phenylindole (DAPI, Thermo Fisher Scientific) to visualize the nuclei. Images were acquired using an EVOS FL Auto 2 Imaging System, using 10x, 20x and 40x air objectives. Brightness and contrast were adjusted identically to the corresponding controls using the Fiji software packages^76^ and figures were assembled using FigureJ^77^. Segmentation of individual cells was based on the ZO-1 staining and performed using CellPose^78^. Outlines were imported and overlayed in Fiji.

#### Cytotox and Cellviabilty Assay

Cytotoxicity in Huh7 knock-out cell lines and upon inhibitor treatment of Huh7 and VeroE6 cell lines was monitored using CytoTox 96^®^ Non-Radioactive Cytotoxicity Assay (Promega, G1780). Relative cytotoxicity compared to lysed control cells was analyzed. Cell viability of primary human nasal epithelial cells was analyzed during inhibitor only treatment at highest concentrations (50 uM, 60 uM) using the CellTiter-Glo^®^ 2.0 Cell Viability Assay (Promega, G9241) and related to DMSO treated cells.

#### LC3-GFP Autophagy

Autophagosome formation was assessed in native Huh7 and Huh7-KO cell lines. Huh7, TMEM41B-KO, MINAR1-KO and FKBP8-KO cells were seeded in a 96 well formation (1.5 Mio cells per plate). LC3-GFP was transfected using Lipofectamine 2000 for 24 hrs. After 24 hrs cells were treated with 100 nM Rapamycin (Sigma Aldrich, S-015) or an equal volume of DMSO for 6 hrs and GFP was analyzed using an EVOS FL Auto 2 Imaging System, using 10x and processed as mentioned above. Alternatively, transfected cells were infected with HCoV-229E at a MOI 0.1 for 24 hrs and GFP expression was analyzed. Images were quantified for autophagosome formation by manual counting using 5 images per condition and three replicates in Fiji. Autophagosome formation was normalized to number of transfected cells.

### Quantification and Statistical Analysis

#### Genome-wide CRISPR/Cas9-mediated KO Screen

For the CRISPR screens, positive enrichment scores, RRA p-values, log fold change (LFC), and false discovery rates were calculated using the MAGeCK algorithm. In Figure S1B, the mean normalized sgRNA counts for each biological replicate were used as input to calculate pairwise correlation. The correlation matrix was generated using the ‘cor’ function in R with the Pearson correlation method and visualized using pheatmap with the clustering performed using correlation as distance metrics.

#### Characterization and Analyses of Top Scoring Host Dependency Factors

Significant difference in data was tested using Nev 2020, version 9.0 or GraphPad Prism version 8.3.1 for Windows (GraphPad). Please refer to figure captions for details regarding the statistical tests applied. *P* values < 0.05 were considered significant.

#### Additional Resources

No additional resources have been created during this study.

