## Supplement for "A genome-wide CRISPR screen identifies interactors of the autophagy pathway as conserved coronavirus targets"

### Figure 1

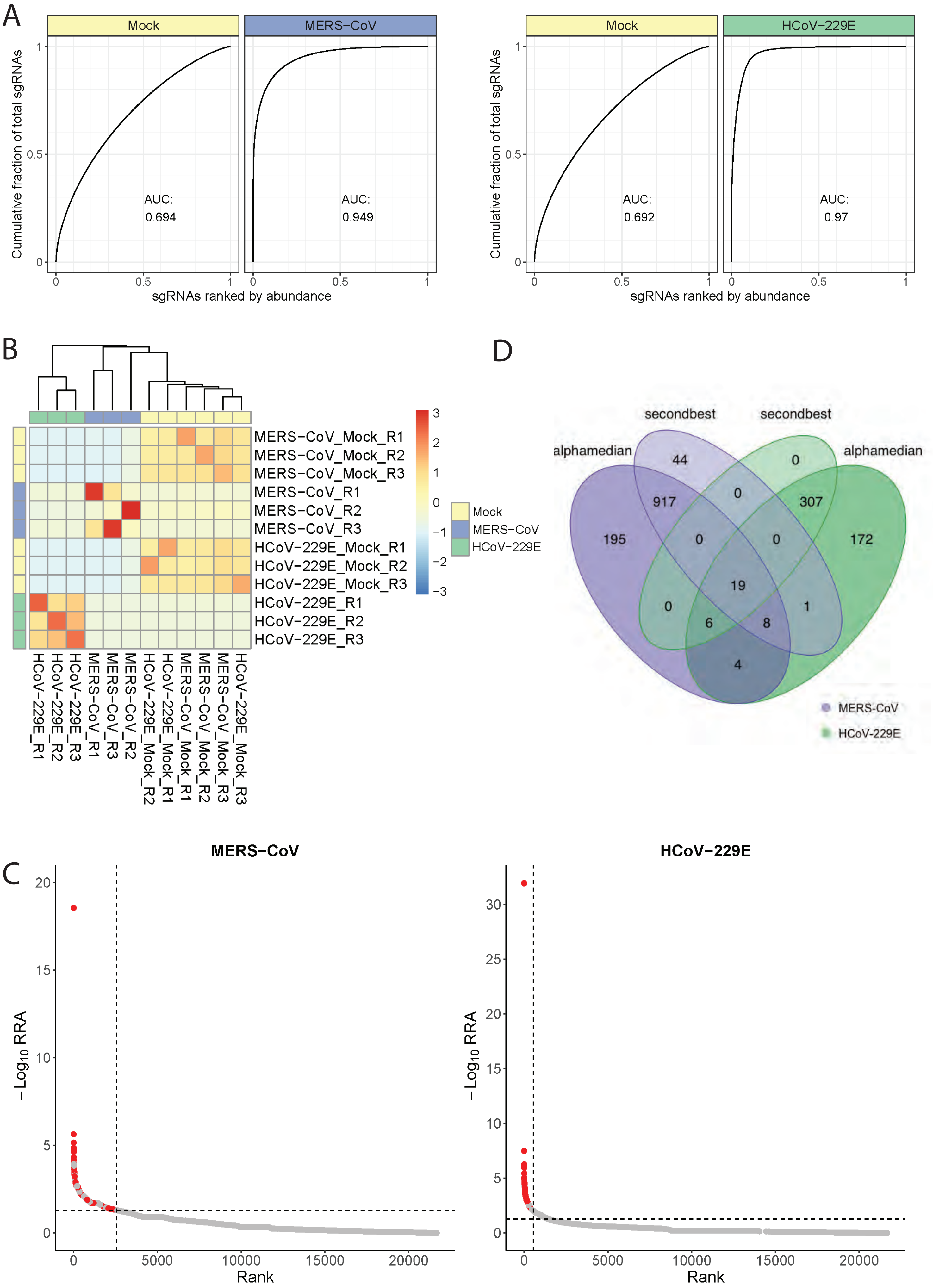
A
Mock
MERS−CoV
Mock
HCoV−229E
Cumulative fraction of total sgRNAs
AUC:
0.694
AUC:
0.949
AUC:
0.692
AUC:
0.97
sgRNAs ranked by abundance
B
MERS−CoV_Mock_R1
MERS−CoV_Mock_R2
MERS−CoV_Mock_R3
MERS−CoV_R1
MERS−CoV_R2
MERS−CoV_R3
HCoV−229E_Mock_R1
HCoV−229E_Mock_R2
HCoV−229E_Mock_R3
HCoV−229E_R1
HCoV−229E_R2
HCoV−229E_R3
Mock
MERS−CoV
HCoV−229E
D
secondbest
secondbest
alphamedian
alphamedian
MERS-CoV
HCoV-229E
C
MERS−CoV
HCoV−229E
−Log10 RRA
Rank

Figure 2

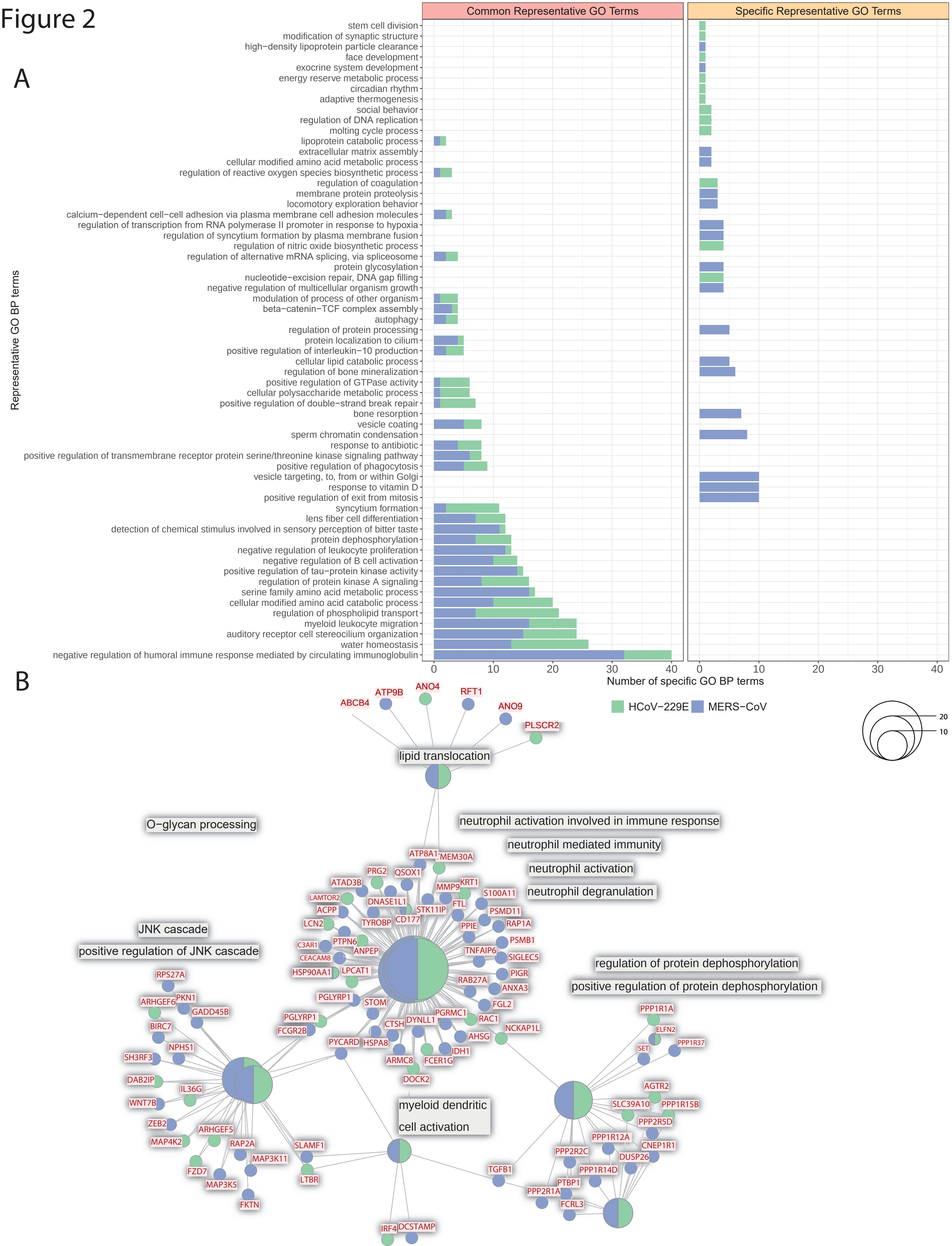

A
Common Representative GO Terms
Specific Representative GO Terms
Representative GO BP terms
stem cell division
modification of synaptic structure
high−density lipoprotein particle clearance
face development
exocrine system development
energy reserve metabolic process
circadian rhythm
adaptive thermogenesis
social behavior
regulation of DNA replication
molting cycle process
lipoprotein catabolic process
extracellular matrix assembly
cellular modified amino acid metabolic process
regulation of reactive oxygen species biosynthetic process
regulation of coagulation
membrane protein proteolysis
locomotory exploration behavior
calcium−dependent cell−cell adhesion via plasma membrane cell adhesion molecules
regulation of transcription from RNA polymerase II promoter in response to hypoxia
regulation of syncytium formation by plasma membrane fusion
regulation of nitric oxide biosynthetic process
regulation of alternative mRNA splicing, via spliceosome
protein glycosylation
nucleotide−excision repair, DNA gap filling
negative regulation of multicellular organism growth
modulation of process of other organism
beta−catenin−TCF complex assembly
autophagy
regulation of protein processing
protein localization to cilium
positive regulation of interleukin−10 production
cellular lipid catabolic process
regulation of bone mineralization
positive regulation of GTPase activity
cellular polysaccharide metabolic process
positive regulation of double−strand break repair
bone resorption
vesicle coating
sperm chromatin condensation
response to antibiotic
positive regulation of transmembrane receptor protein serine/threonine kinase signaling pathway
positive regulation of phagocytosis
vesicle targeting, to, from or within Golgi
response to vitamin D
positive regulation of exit from mitosis
syncytium formation
lens fiber cell differentiation
detection of chemical stimulus involved in sensory perception of bitter taste
protein dephosphorylation
negative regulation of leukocyte proliferation
negative regulation of B cell activation
positive regulation of tau−protein kinase activity
regulation of protein kinase A signaling
serine family amino acid metabolic process
cellular modified amino acid catabolic process
regulation of phospholipid transport
myeloid leukocyte migration
auditory receptor cell stereocilium organization
water homeostasis
negative regulation of humoral immune response mediated by circulating immunoglobulin
0 10 20 30 40
0 10 20 30 40
Number of specific GO BP terms
HCoV−229E
MERS−CoV
B
20
10
lipid translocation
ABCB4
ATP9B
ANO4
RFT1
ANO9
PLSCR2
O−glycan processing
neutrophil activation involved in immune response
neutrophil mediated immunity
neutrophil activation
neutrophil degranulation
JNK cascade
positive regulation of JNK cascade
regulation of protein dephosphorylation
positive regulation of protein dephosphorylation
myeloid dendritic cell activation
ATP8A1
TMEM30A
PRG2
QSOX1
ATAD3B
KRT1
MMP9
S100A11
LAMTOR2
DNASE1L1
FTL
PSMD11
ACPP
STK11IP
LCN2
TYROBP
CD177
PPIE
RAP1A
PSMB1
C3AR1
PTPN6
TNFAIP6
SIGLEC5
CEACAM8
ANPEP
PIGR
HSP90AA1
LPCAT1
RAB27A
ANXA3
PGLYRP1
STOM
FGL2
PGRMC1
RAC1
PGLYRP1
DYNLL1
FCGR2B
CTSH
AHSG
NCKAP1L
PYCARD
HSPA8
IDH1
ARMC8
FCER1G
DOCK2
RPS27A
PKN1
ARHGEF6
GADD45B
BIRC7
NPHS1
SH3RF3
DAB2IP
IL36G
WNT7B
ZEB2
ARHGEF5
MAP4K2
RAP2A
SLAMF1
MAP3K11
FZD7
LTBR
MAP3K5
FKTN
IRF4
DCSTAMP
TGFB1
PPP1R1A
ELFN2
PPP1R37
SET
AGTR2
SLC39A10
PPP1R15B
PPP2R5D
PPP1R12A
CNEP1R1
PPP2R2C
DUSP26
PPP1R14D
PTBP1
PPP2R1A
FCRL3

Figure 3

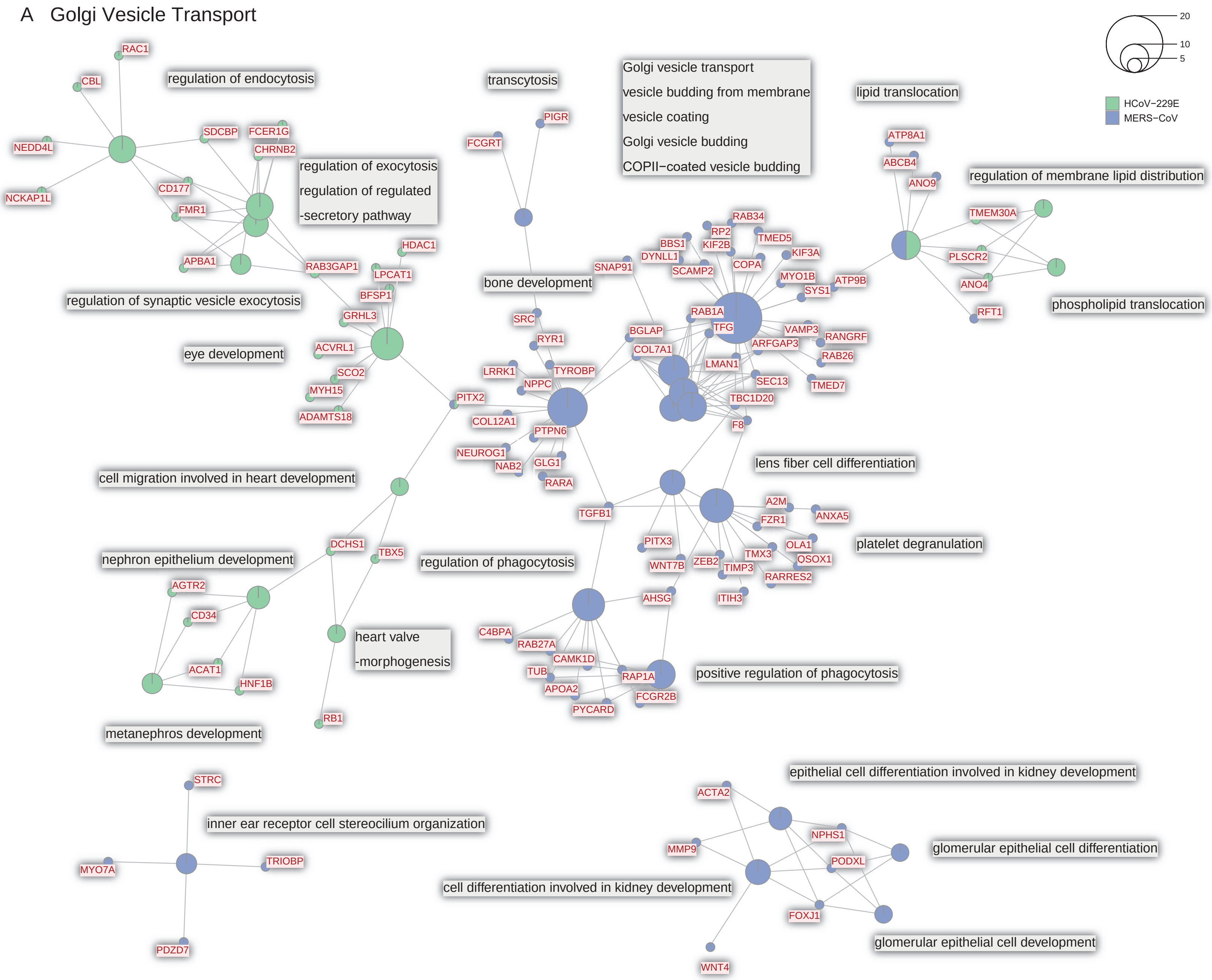

A Golgi Vesicle Transport
20
10
5
HCoV−229E
MERS−CoV
regulation of endocytosis
RAC1
CBL
NEDD4L
NCKAP1L
SDCBP
FCER1G
CHRNB2
CD177
FMR1
APBA1
RAB3GAP1
HDAC1
LPCAT1
BFSP1
GRHL3
ACVRL1
SCO2
MYH15
ADAMTS18
regulation of exocytosis
regulation of regulated
-secretory pathway
regulation of synaptic vesicle exocytosis
eye development
transcytosis
PIGR
FCGRT
bone development
SRC
RYR1
TYROBP
LRRK1
NPPC
PITX2
COL12A1
PTPN6
NEUROG1
NAB2
GLG1
RARA
TGFB1
Golgi vesicle transport
vesicle budding from membrane
vesicle coating
Golgi vesicle budding
COPII−coated vesicle budding
SNAP91
BBS1
DYNLL1
RAB34
RP2
KIF2B
TMED5
SCAMP2
COPA
KIF3A
MYO1B
SYS1
ATP9B
RAB1A
TFG
BGLAP
COL7A1
VAMP3
ARFGAP3
RANGRF
RAB26
LMAN1
SEC13
TMED7
TBC1D20
F8
lipid translocation
ATP8A1
ABCB4
ANO9
regulation of membrane lipid distribution
TMEM30A
PLSCR2
ANO4
RFT1
phospholipid translocation
lens fiber cell differentiation
A2M
ANXA5
FZR1
platelet degranulation
PITX3
OLA1
TMX3
OSOX1
WNT7B
ZEB2
TIMP3
RARRES2
AHSG
ITIH3
regulation of phagocytosis
C4BPA
RAB27A
CAMK1D
TUB
APOA2
RAP1A
FCGR2B
PYCARD
positive regulation of phagocytosis
cell migration involved in heart development
DCHS1
TBX5
nephron epithelium development
AGTR2
CD34
ACAT1
HNF1B
heart valve
-morphogenesis
RB1
metanephros development
STRC
inner ear receptor cell stereocilium organization
TRIOBP
MYO7A
PDZD7
epithelial cell differentiation involved in kidney development
ACTA2
NPHS1
glomerular epithelial cell differentiation
MMP9
PODXL
cell differentiation involved in kidney development
FOXJ1
glomerular epithelial cell development
WNT4

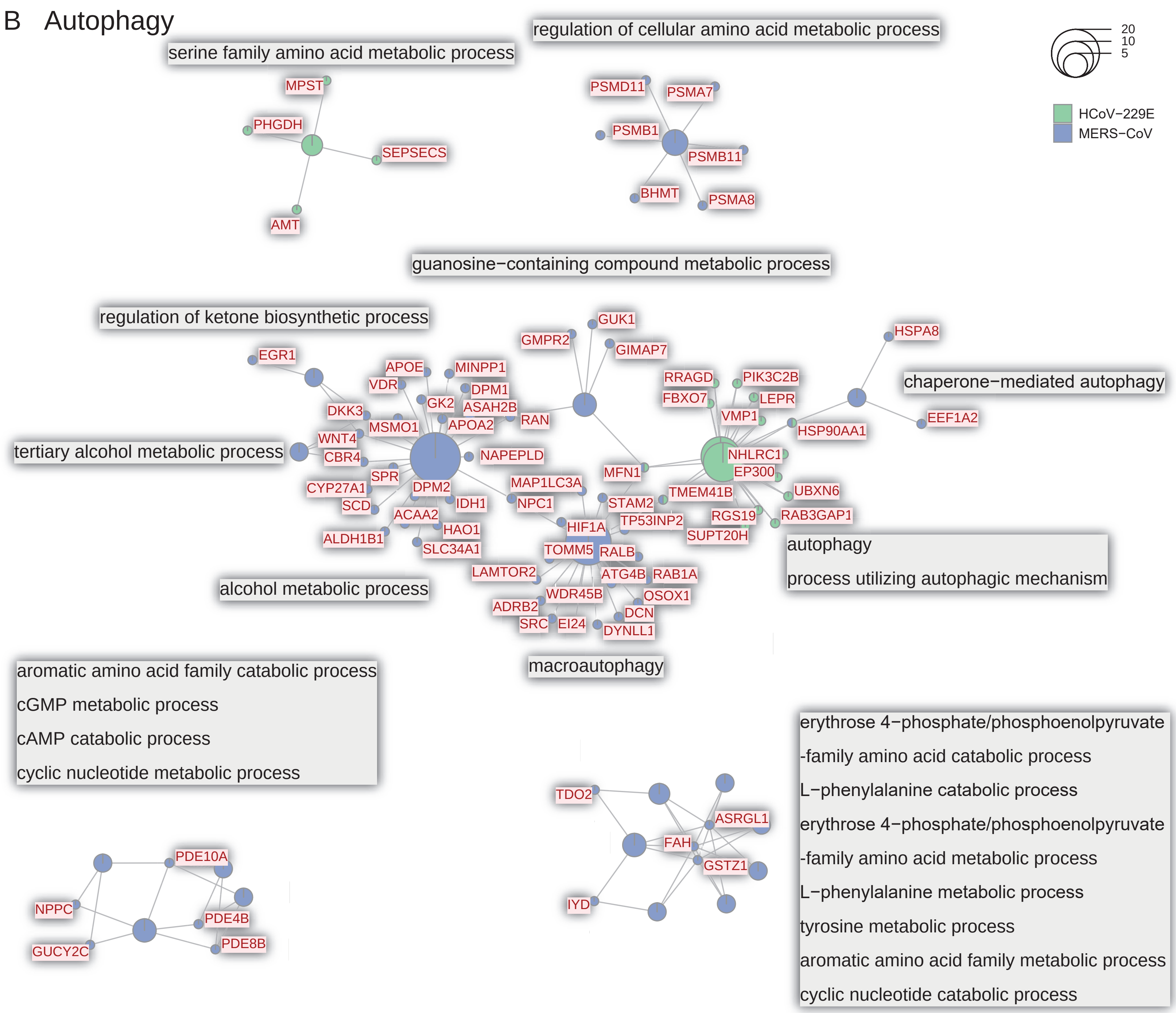
B Autophagy
serine family amino acid metabolic process
MPST
PHGDH
SEPSECS
AMT
regulation of cellular amino acid metabolic process
PSMD11
PSMA7
PSMB1
PSMB11
BHMT
PSMA8
20
10
5
HCoV−229E
MERS−CoV
guanosine−containing compound metabolic process
regulation of ketone biosynthetic process
GUK1
GMPR2
GIMAP7
HSPA8
EGR1
APOE
MINPP1
VDR
DPM1
GK2
ASAH2B
DKK3
MSMO1
APOA2
RAN
RRAGD
FBXO7
PIK3C2B
LEPR
VMP1
chaperone−mediated autophagy
EEF1A2
HSP90AA1
tertiary alcohol metabolic process
WNT4
CBR4
NAPEPLD
SPR
CYP27A1
DPM2
SCD
IDH1
ACAA2
HAO1
ALDH1B1
SLC34A1
MAP1LC3A
NPC1
MFN1
NHLRC1
EP300
TMEM41B
UBXN6
STAM2
TP53INP2
RGS19
RAB3GAP1
SUPT20H
HIF1A
TOMM5
RALB
autophagy
process utilizing autophagic mechanism
LAMTOR2
ATG4B
RAB1A
alcohol metabolic process
WDR45B
OSOX1
ADRB2
DCN
SRC
EI24
DYNLL1
macroautophagy
aromatic amino acid family catabolic process
cGMP metabolic process
cAMP catabolic process
cyclic nucleotide metabolic process
erythrose 4−phosphate/phosphoenolpyruvate
-family amino acid catabolic process
L−phenylalanine catabolic process
erythrose 4−phosphate/phosphoenolpyruvate
-family amino acid metabolic process
L−phenylalanine metabolic process
tyrosine metabolic process
aromatic amino acid family metabolic process
cyclic nucleotide catabolic process
TDO2
ASRGL1
FAH
GSTZ1
IYD
PDE10A
NPPC
PDE4B
GUCY2C
PDE8B

C Catabolic Processes

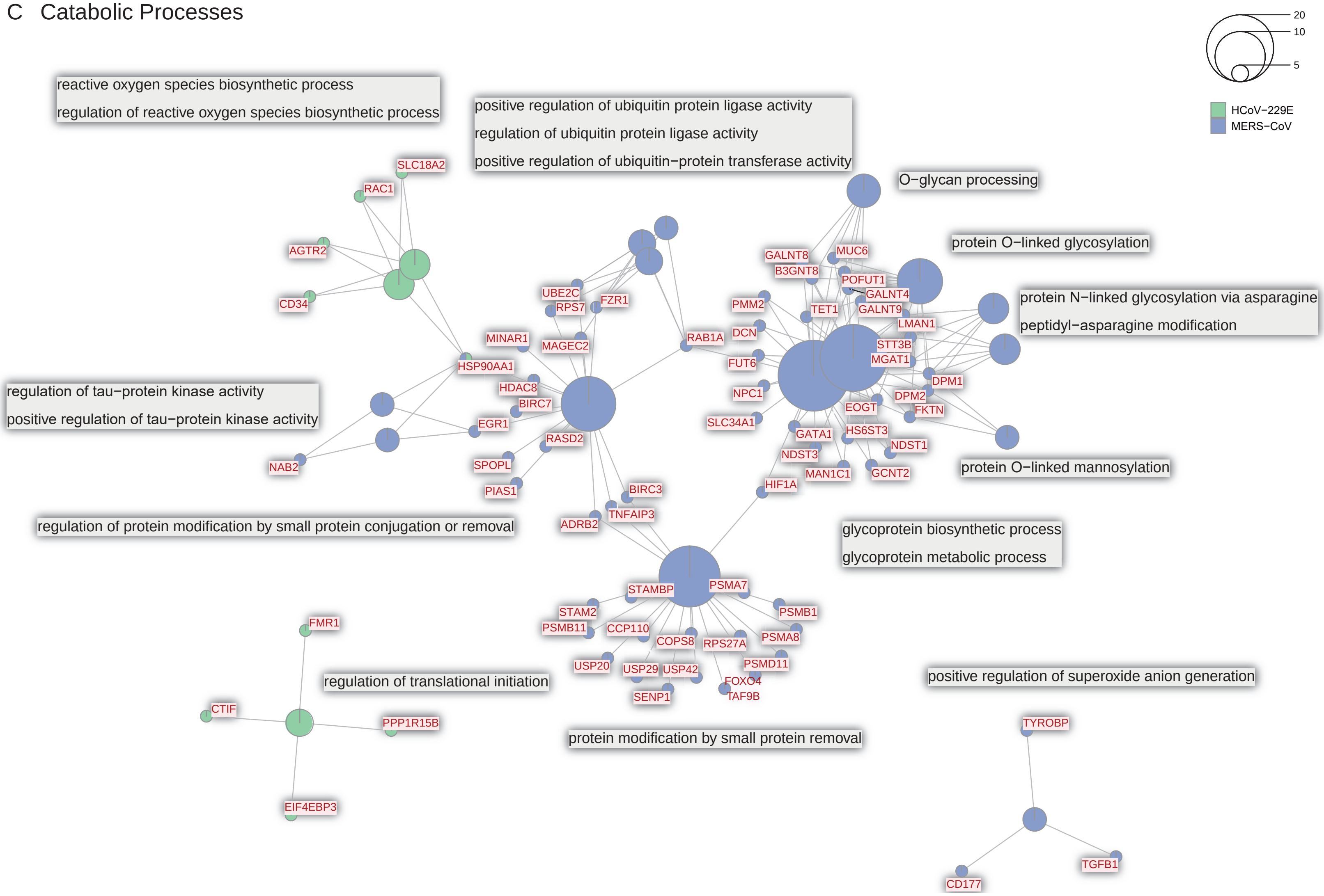

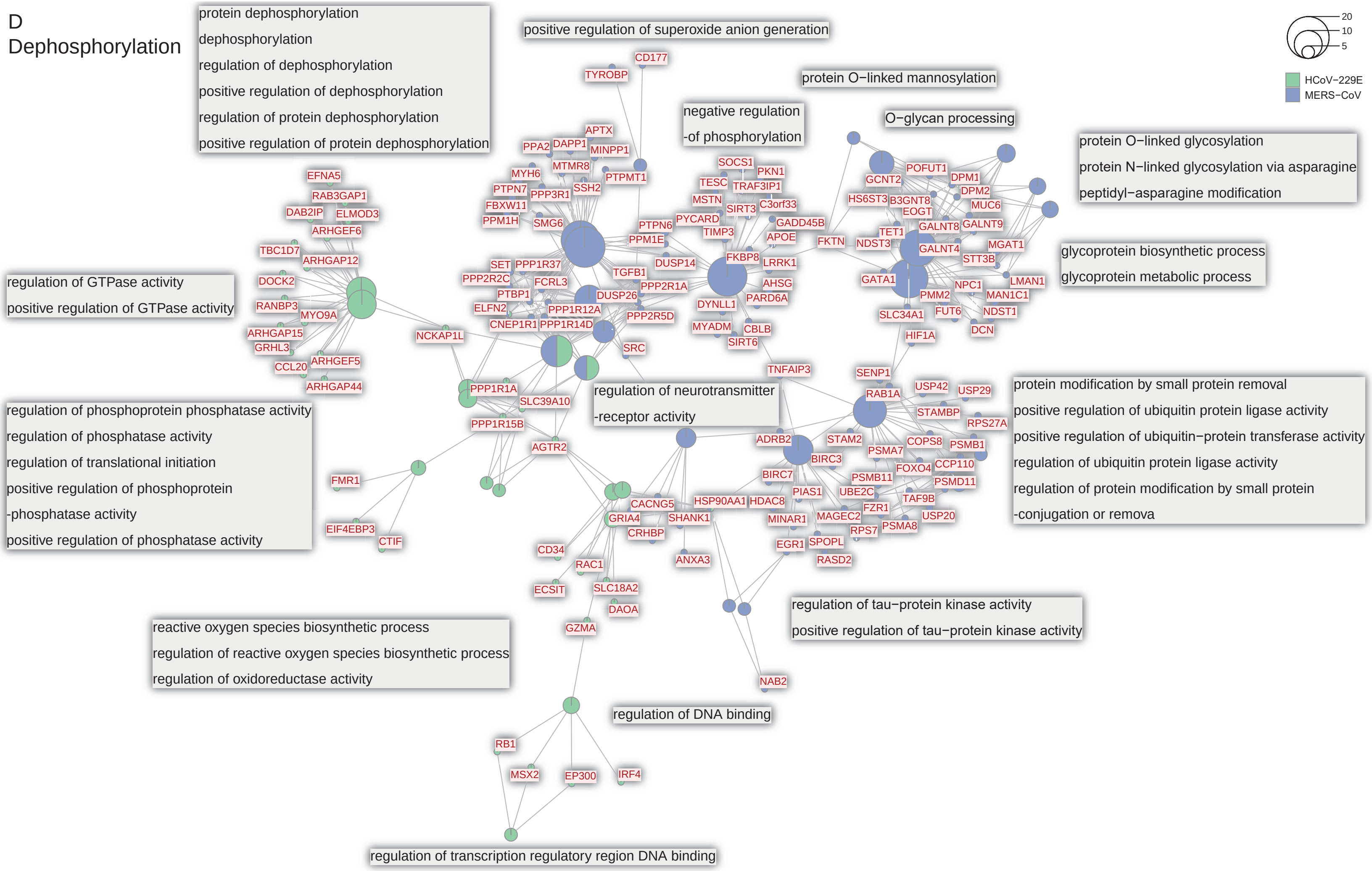
D
Dephosphorylation
protein dephosphorylation
dephosphorylation
regulation of dephosphorylation
positive regulation of dephosphorylation
regulation of protein dephosphorylation
positive regulation of protein dephosphorylation
positive regulation of superoxide anion generation
protein O−linked mannosylation
negative regulation
-of phosphorylation
O−glycan processing
protein O−linked glycosylation
protein N−linked glycosylation via asparagine
peptidyl−asparagine modification
glycoprotein biosynthetic process
glycoprotein metabolic process
regulation of GTPase activity
positive regulation of GTPase activity
regulation of phosphoprotein phosphatase activity
regulation of phosphatase activity
regulation of translational initiation
positive regulation of phosphoprotein
-phosphatase activity
positive regulation of phosphatase activity
regulation of neurotransmitter
-receptor activity
protein modification by small protein removal
positive regulation of ubiquitin protein ligase activity
positive regulation of ubiquitin−protein transferase activity
regulation of ubiquitin protein ligase activity
regulation of protein modification by small protein
-conjugation or remova
regulation of tau−protein kinase activity
positive regulation of tau−protein kinase activity
reactive oxygen species biosynthetic process
regulation of reactive oxygen species biosynthetic process
regulation of oxidoreductase activity
regulation of DNA binding
regulation of transcription regulatory region DNA binding
20
10
5
HCoV−229E
MERS−CoV
CD177
TYROBP
APTX
PPA2
DAPP1
MINPP1
MYH6
MTMR8
PTPMT1
PTPN7
PPP3R1
SSH2
FBXW11
PPM1H
SMG6
PTPN6
PPM1E
SOCS1
PKN1
TESC
TRAF3IP1
MSTN
SIRT3
C3orf33
PYCARD
GADD45B
TIMP3
APOE
FKBP8
LRRK1
DUSP14
AHSG
PARD6A
DYNLL1
MYADM
CBLB
SIRT6
GCNT2
POFUT1
DPM1
HS6ST3
B3GNT8
DPM2
EOGT
MUC6
TET1
GALNT8
GALNT9
FKTN
NDST3
GALNT4
MGAT1
STT3B
GATA1
NPC1
LMAN1
PMM2
MAN1C1
FUT6
NDST1
SLC34A1
DCN
HIF1A
EFNA5
RAB3GAP1
DAB2IP
ELMOD3
ARHGEF6
TBC1D7
ARHGAP12
DOCK2
RANBP3
MYO9A
ARHGAP15
GRHL3
CCL20
ARHGEF5
ARHGAP44
SET
PPP1R37
PPP2R2C
FCRL3
TGFB1
PPP2R1A
PTBP1
DUSP26
ELFN2
PPP1R12A
PPP2R5D
CNEP1R1
PPP1R14D
NCKAP1L
SRC
PPP1R1A
SLC39A10
PPP1R15B
AGTR2
FMR1
EIF4EBP3
CTIF
TNFAIP3
SENP1
USP42
USP29
RAB1A
STAMBP
RPS27A
ADRB2
STAM2
PSMA7
COPS8
PSMB1
BIRC3
FOXO4
CCP110
BIRC7
PSMB11
PSMD11
PIAS1
UBE2C
TAF9B
HSP90AA1
HDAC8
FZR1
CACNG5
MINAR1
MAGEC2
USP20
GRIA4
SHANK1
RPS7
PSMA8
CRHBP
EGR1
SPOPL
CD34
RASD2
ANXA3
RAC1
ECSIT
SLC18A2
DAOA
GZMA
NAB2
RB1
MSX2
EP300
IRF4

E Immunity

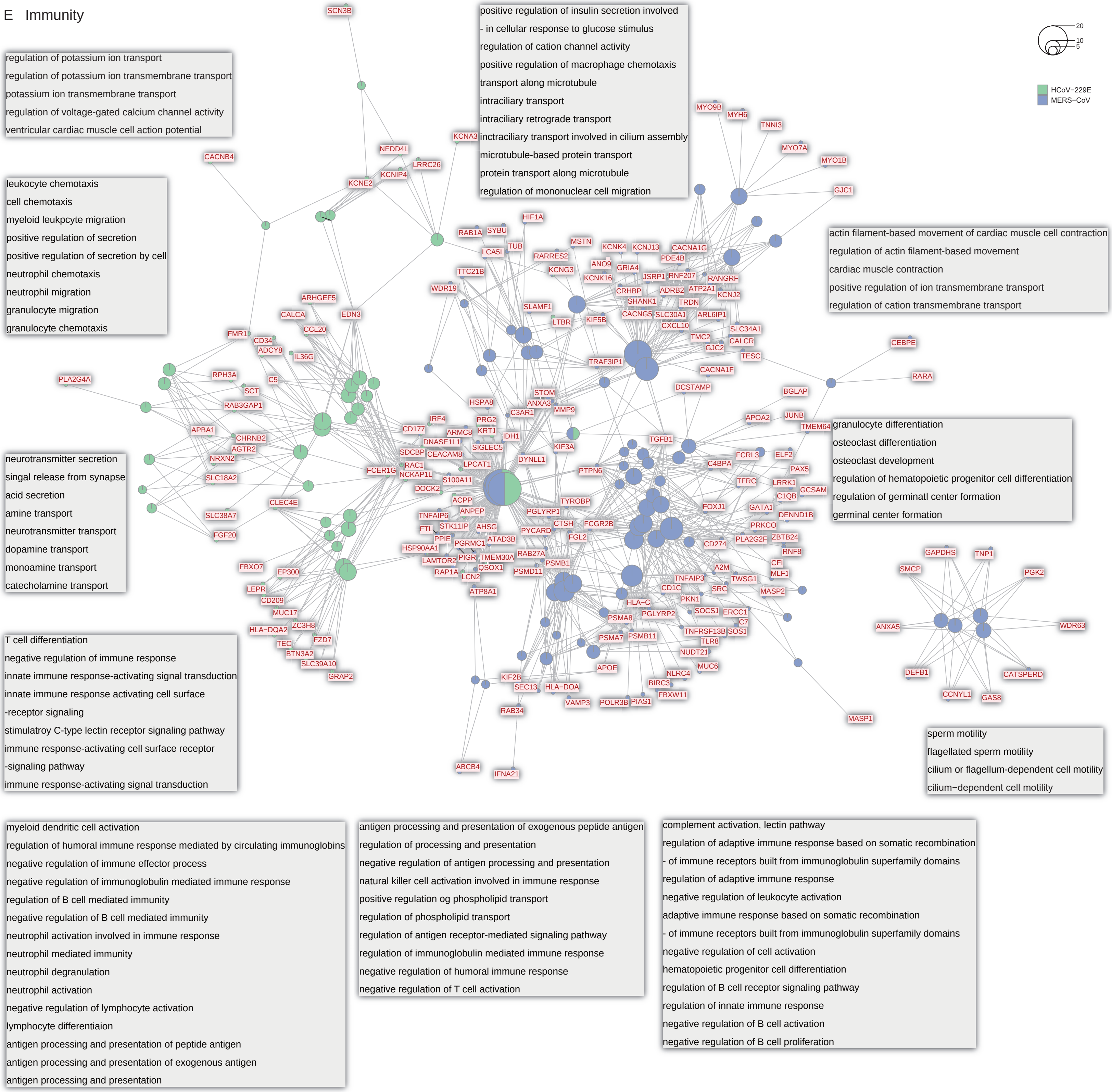

regulation of potassium ion transport
regulation of potassium ion transmembrane transport
potassium ion transmembrane transport
regulation of voltage-gated calcium channel activity
ventricular cardiac muscle cell action potential
positive regulation of insulin secretion involved
- in cellular response to glucose stimulus
regulation of cation channel activity
positive regulation of macrophage chemotaxis
transport along microtubule
intraciliary transport
intraciliary retrograde transport
inctraciliary transport involved in cilium assembly
microtubule-based protein transport
protein transport along microtubule
regulation of mononuclear cell migration
20
10
5
HCoV−229E
MERS−CoV
leukocyte chemotaxis
cell chemotaxis
myeloid leukpcyte migration
positive regulation of secretion
positive regulation of secretion by cell
neutrophil chemotaxis
neutrophil migration
granulocyte migration
granulocyte chemotaxis
actin filament-based movement of cardiac muscle cell contraction
regulation of actin filament-based movement
cardiac muscle contraction
positive regulation of ion transmembrane transport
regulation of cation transmembrane transport
neurotransmitter secretion
singal release from synapse
acid secretion
amine transport
neurotransmitter transport
dopamine transport
monoamine transport
catecholamine transport
granulocyte differentiation
osteoclast differentiation
osteoclast development
regulation of hematopoietic progenitor cell differentiation
regulation of germinatl center formation
germinal center formation
T cell differentiation
negative regulation of immune response
innate immune response-activating signal transduction
innate immune response activating cell surface
-receptor signaling
stimulatroy C-type lectin receptor signaling pathway
immune response-activating cell surface receptor
-signaling pathway
immune response-activating signal transduction
sperm motility
flagellated sperm motility
cilium or flagellum-dependent cell motility
cilium−dependent cell motility
myeloid dendritic cell activation
regulation of humoral immune response mediated by circulating immunoglobins
negative regulation of immune effector process
negative regulation of immunoglobulin mediated immune response
regulation of B cell mediated immunity
negative regulation of B cell mediated immunity
neutrophil activation involved in immune response
neutrophil mediated immunity
neutrophil degranulation
neutrophil activation
negative regulation of lymphocyte activation
lymphocyte differentiaion
antigen processing and presentation of peptide antigen
antigen processing and presentation of exogenous antigen
antigen processing and presentation
antigen processing and presentation of exogenous peptide antigen
regulation of processing and presentation
negative regulation of antigen processing and presentation
natural killer cell activation involved in immune response
positive regulation og phospholipid transport
regulation of phospholipid transport
regulation of antigen receptor-mediated signaling pathway
regulation of immunoglobulin mediated immune response
negative regulation of humoral immune response
negative regulation of T cell activation
complement activation, lectin pathway
regulation of adaptive immune response based on somatic recombination
- of immune receptors built from immunoglobulin superfamily domains
regulation of adaptive immune response
negative regulation of leukocyte activation
adaptive immune response based on somatic recombination
- of immune receptors built from immunoglobulin superfamily domains
negative regulation of cell activation
hematopoietic progenitor cell differentiation
regulation of B cell receptor signaling pathway
regulation of innate immune response
negative regulation of B cell activation
negative regulation of B cell proliferation

F Developmental Processes

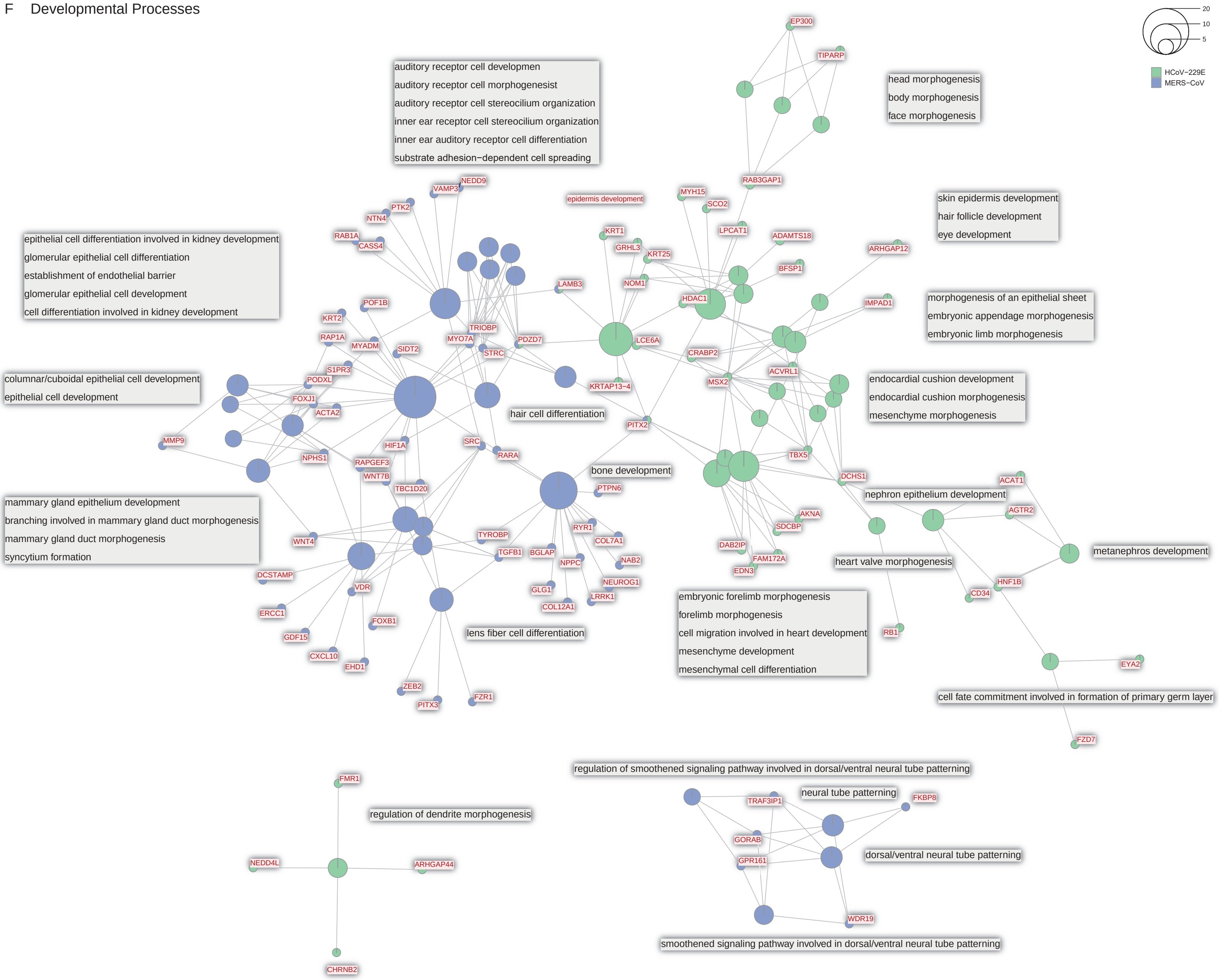

G Homeostatic Processes

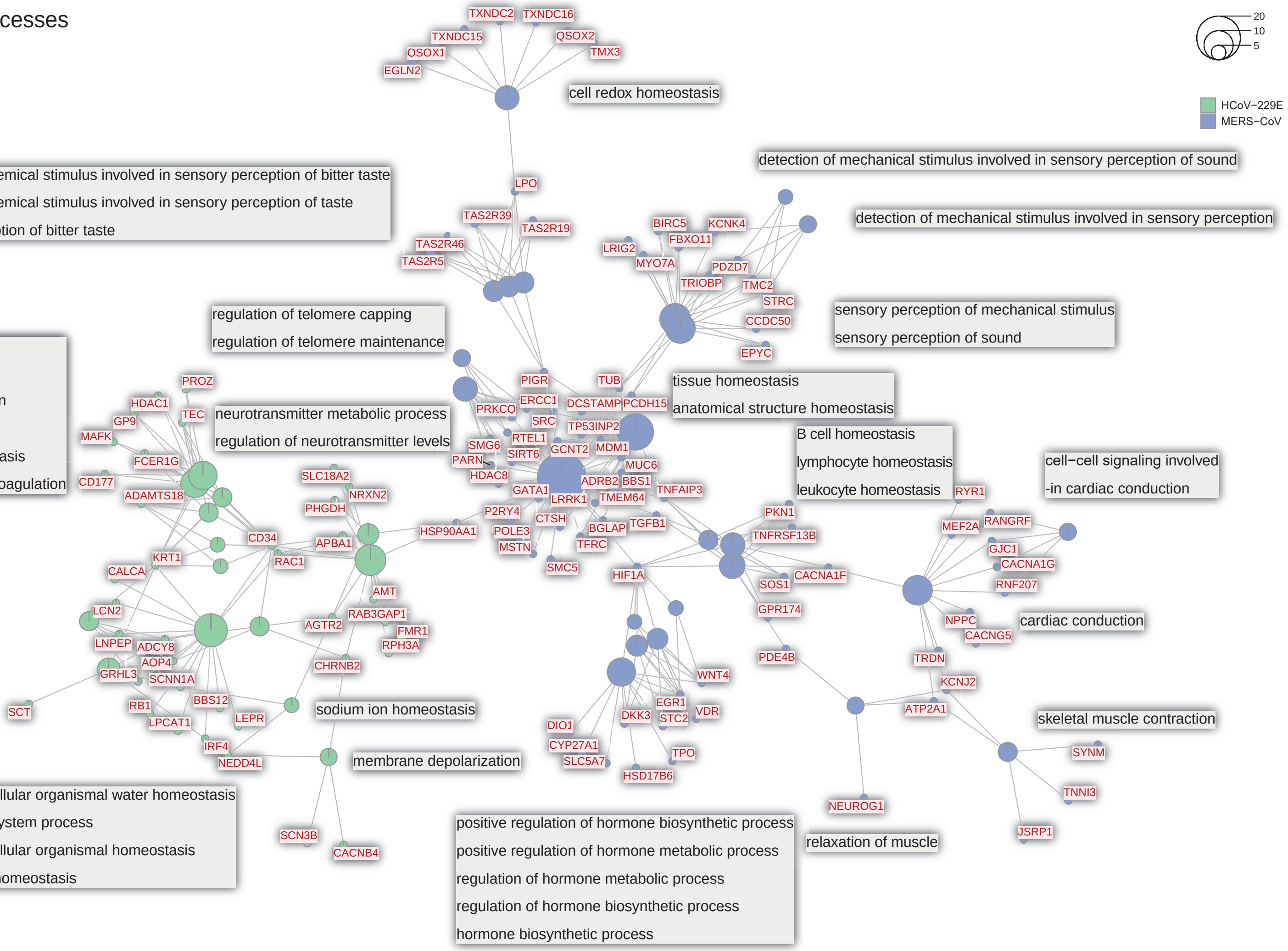

Figure 4

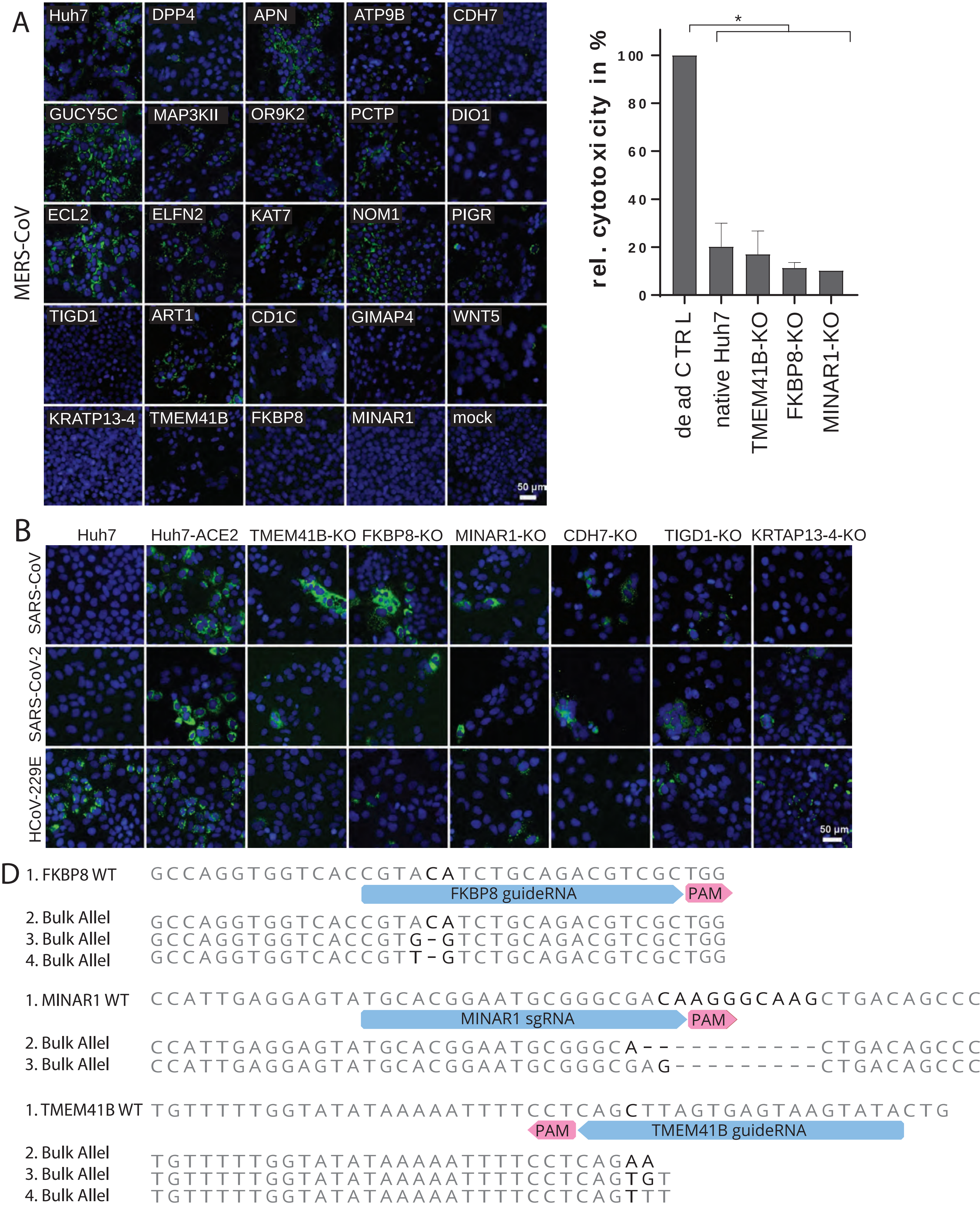
A
MERS-CoV
Huh7
DPP4
APN
ATP9B
CDH7
GUCY5C
MAP3KII
OR9K2
PCTP
DIO1
ECL2
ELFN2
KAT7
NOM1
PIGR
TIGD1
ART1
CD1C
GIMAP4
WNT5
KRATP13-4
TMEM41B
FKBP8
MINAR1
mock
50 μm
C
rel. cytotoxicity in %
0
20
40
60
80
100
*
dead CTRL
native Huh7
TMEM41B-KO
FKBP8-KO
MINAR1-KO
B
Huh7
Huh7-ACE2
TMEM41B-KO
FKBP8-KO
MINAR1-KO
CDH7-KO
TIGD1-KO
KRTAP13-4-KO
SARS-CoV
SARS-CoV-2
HCoV-229E
50 μm
D
1. FKBP8 WT GCCAGGTGGTCACCGTACATCTGCAGACGTCGCTGG
FKBP8 guideRNA
PAM
2. Bulk Allel GCCAGGTGGTCACCGTACATCTGCAGACGTCGCTGG
3. Bulk Allel GCCAGGTGGTCACCGTG-GTCTGCAGACGTCGCTGG
4. Bulk Allel GCCAGGTGGTCACCGTT-GTCTGCAGACGTCGCTGG
1. MINAR1 WT CCATTGAGGAGTATGCACGGAATGCGGGCGACAAGGGCAAGCTGACAGCCC
MINAR1 sgRNA
PAM
2. Bulk Allel CCATTGAGGAGTATGCACGGAATGCGGGCA-----------CTGACAGCCC
3. Bulk Allel CCATTGAGGAGTATGCACGGAATGCGGGCGAG----------CTGACAGCCC
1. TMEM41B WT TGTTTTTGGTATATAAAAATTTTCCTCAGCTTAGTGAGTAAGTATACTG
PAM
TMEM41B guideRNA
2. Bulk Allel TGTTTTTGGTATATAAAAATTTTCCTCAGAA
3. Bulk Allel TGTTTTTGGTATATAAAAATTTTCCTCAGTGT
4. Bulk Allel TGTTTTTGGTATATAAAAATTTTCCTCAGTTT

### Figure 5

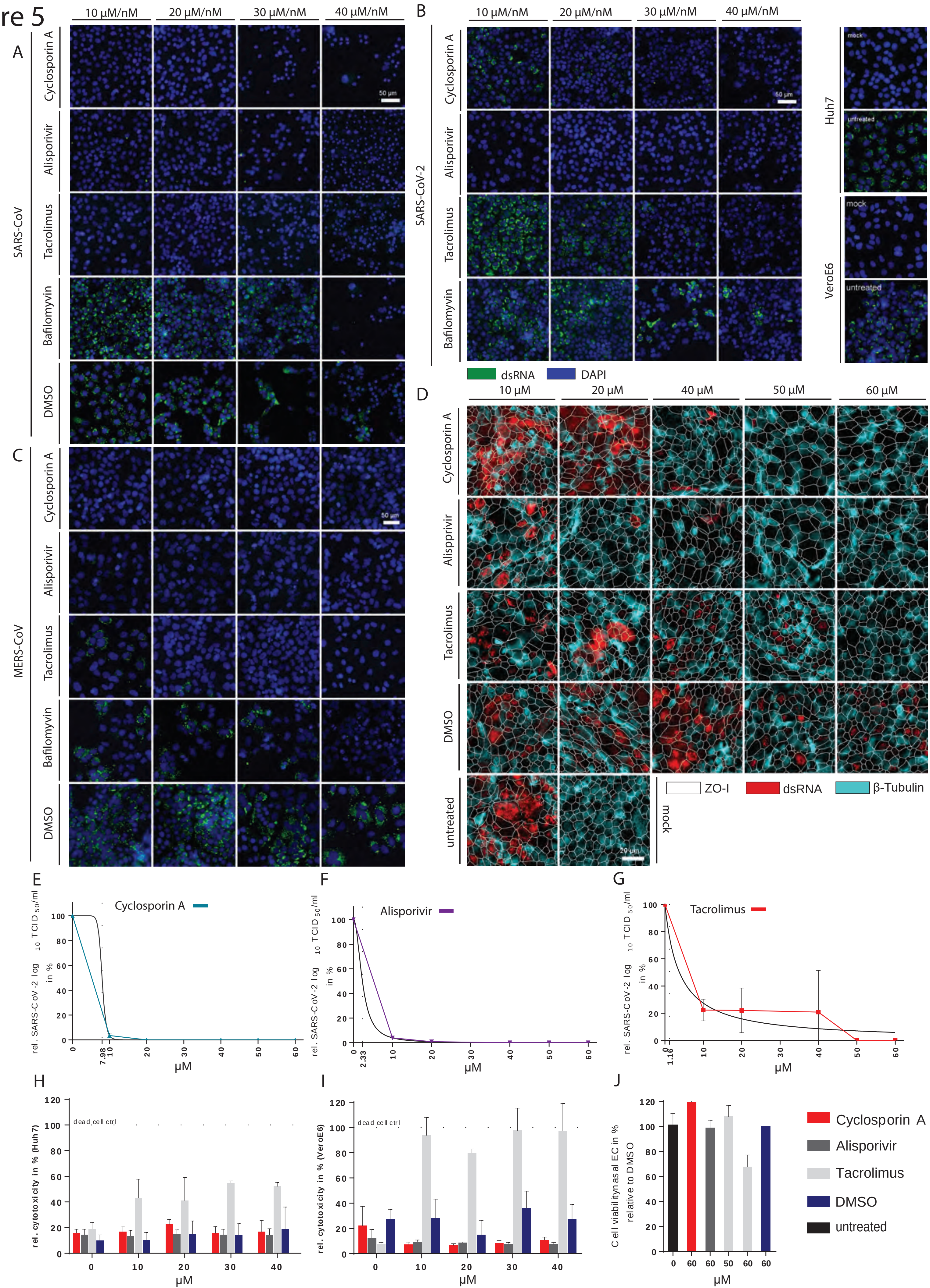

A
B
C
D
E
F
G
H
I
J
10 µM/nM
20 µM/nM
30 µM/nM
40 µM/nM
SARS-CoV
SARS-CoV-2
MERS-CoV
Cyclosporin A
Alisporivir
Tacrolimus
Bafilomyvin
DMSO
50 µm
Huh7
VeroE6
mock
untreated
dsRNA
DAPI
10 µM
20 µM
40 µM
50 µM
60 µM
Alispprivir
ZO-I
β-Tubulin
20 µm
rel. SARS-CoV-2 log 10 TCID 50/ml in %
µM
7.98
2.33
1.16
rel. cytotoxicity in % (Huh7)
rel. cytotoxicity in % (VeroE6)
dead cell ctrl
Cell viability nasal EC in % relative to DMSO

**Supplemental figure titles and legends**

**Figure S1: Quality control metrics and enriched gene identification for MERS-CoV and HCoV-229E genome-wide CRISPR screens.** (A) Area under the curve (AUC) analysis of MERS-CoV and HCoV-229E CRISPR screens evaluating sgRNA library representation in surviving Huh7 cells from uninfected (Mock) and MERS-CoV (left two panels) or HCoV-229E (right two panels) infected samples. For each CRISPR screen, sgRNA abundance was calculated based on average sgRNA abundance over 3 independent biological replicates. (B) Correlation matrix depicting the Pearson correlation for guide-level normalized read counts among biological replicates and samples from both screens. R1, R2, and R3 represent the biological replicates 1, 2, and 3, respectively. Clustering was performed in pheatmap using correlation as a distance metric (C) Robust Rank Aggregation (RRA) p-value distribution of all genes in the GeCKOv2 library for both MERS-CoV (left) and HCoV-229E (right) CRISPR screens. Genes that met the criteria for significance (RRA p-value $\leq 0.05$ and FC $\geq 2$) are highlighted in red. (D) Venn diagram illustrating the overlap between significantly enriched genes from both CRISPR screens that were identified via two different RRA-based analysis methods (alpha median and second best). A total of 19 genes were identified by both methods in both MERS-CoV and HCoV-229E CRISPR screens.

**Figure S2**: (A) Representative GO terms identified using full list of enriched GO terms for MERS-CoV and HCoV-229E screens (Table 2). Representative terms found in both screens are shown in the top panel, whereas virus-specific terms are shown in the bottom panel. BP, CC, and MF represent different GO term categories. (B) Specific GO terms enriched in both CoV screens (individual GO terms, not representative GO terms).

**Figure S3:** Cnet plots for GO BP terms found in each individual biological cluster shown in Figure 2A (S3A Golgi Vesicle Transport, S3B Autophagy, S3C Catabolic Processes, S3D Dephosphorylation, S3E Immunity, S3F Developmental Processes, S3G Homeostatic Processes). Plots include both GO terms that contain one or more of the 19 common significantly enriched genes found in both CoV screens (as in Figure 2A and 2B) as well as representative GO terms found in both screens that do not contain these genes. Each plot shows the relationship among individual GO terms and genes found in each biological cluster. Larger nodes represent individual GO terms and smaller nodes represent individual gene. Nodes that are functionally related cluster together into a larger network. Node size reflects the number of significantly enriched genes in the node and color indicates the CoV screen for which the node was significant.

**Figure S4: CRISPR-mediated KO of top scoring host dependency factors impairs CoV replication.** (A) Immunofluorescence staining of MERS-CoV infected of Huh7 cells containing KO of top scoring HDFs. dsRNA is shown in green, DAPI is shown in blue. (B) Immunofluorescence staining of HCoV-229E, SARS-CoV and SARS-CoV-2 infected Huh7 cells with TMEM41B, FKBP8 and MINAR1-KO, as well as a stable ACE2 expression. dsRNA is shown in green, DAPI is shown in blue, ACE2 is shown in red. Scale bar is 50 µm. All images were acquired using an Evos Auto FL2 and processed in Fiji. (D) Relative cytotoxicity of TMEM41-KO, FKBP8-KO and MINAR1-KO is depicted in %. Two tailed unpaired student t-test was used to determine significance in GraphPad Prism 8.3.1. (E) Sanger sequencing of FKBP8-KO, MINAR1-KO and TMEM41B-KO verifies Cas9-mediated double strand break in multiple alleles of the KO cells. PAM sequence is indicated in red, binding site of sgRNA is indicated in blue.

**Figure S5: Cyclosporin A, Alisporivir and Tacrolimus inhibit CoV infection in a dose-dependent manner in cell lines and primary human nasal epithelial cells at non-cytotoxic concentrations.** Immunofluorescence staining of SARS-CoV (A), as well as SARS-CoV-2 (B) infected VeroE6 cells and MERS-CoV (C) infected Huh7 cells following Cyclosporin A, Alisporivir, Tacrolimus treatment at 10 µM to 40 µM and Bafilomycin treatment at 10 nM – 40 nM, as well as well as DMSO CTRL as respective volumes 24 hrs post infection/inhibitor treatment. dsRNA is shown in green, DAPI is shown in blue. Scale bar is 50 µm. (D) Immunofluorescence staining of SARS-CoV-2 infected and Cyclosporin A, Alisporivir and Tacrolimus, as well as DMSO treated primary human nasal epithelial cells at 10 µM to 60 µM 48 hpi/post inhibitor treatment. dsRNA (red), cilia (β-Tubulin, light blue) and the outline of segmented cells (ZO-I, white) are shown. Scale bar is 20 µm. All images were acquired using an EVOS FL Auto 2 imaging system with a 10x air objective (A, B, C) and a 40x air objective (D) and processed using Fiji. (E-G) Inhibitor treated primary nasal epithelial cell cultures displayed as inhibitor versus normalized response. $IC_{50}$ value is marked with dotted line and indicated on y axis. Calculations were performed in GraphPad Prism 8.3.1. Cyclosporin A, Alisporivir andTacrolimustreatment-mediated cytotoxicity in Huh7 cells (H) and VeroE6 cells (I) shown relative to dead cell control. (J) Relative nasal epithelial cell culture viability upon treatment of (50 μM and) 60 μM Cyclosporin A, Alisporivir and Tacrolimus normalized to DMSO.
